# Crystal structure of MutYX: A novel clusterless adenine DNA glycosylase with a distinct C-terminal domain and 8-Oxoguanine recognition sphere

**DOI:** 10.1101/2025.01.03.631205

**Authors:** C.H. Trasviña-Arenas, Mohammad Hashemian, Melody Malek, Steven Merrill, Andrew J. Fisher, Sheila S. David

## Abstract

The [4Fe-4S] cluster is an important cofactor of the base excision repair (BER) adenine DNA glycosylase MutY to prevent mutations associated with 8-oxoguanine (OG). Several MutYs lacking the [4Fe-4S] cofactor have been identified. Phylogenetic analysis shows that clusterless MutYs are distributed in two clades suggesting cofactor loss in two independent evolutionary events. Herein, we determined the first crystal structure of a clusterless MutY complexed with DNA. On the basis of the dramatic structural divergence from canonical MutYs, we refer to this as representative of a clusterless MutY subgroup “MutYX”. Interestingly, MutYX compensates for the missing [4Fe-4S] cofactor to maintain positioning of catalytic residues by expanding a pre-existing α-helix and acquisition of the new α-helix. Surprisingly, MutYX also acquired a new C-terminal domain that uniquely recognizes OG using residue Gln201 and Arg209. Adenine glycosylase assays and binding affinity measurements indicate that Arg209 is the primary residue responsible to specificity for OG:A lesions, while Gln201 bridges OG and Arg209. Surprisingly, replacement of Arg209 and Gln201 with Ala increases activity toward G:A mismatches. The MutYX structure serves as an example of devolution, capturing structural features required to retain function in the absence of a metal cofactor considered indispensable.

## INTRODUCTION

Bacteria, archaea and eukaryotes harbor MutY enzymes to prevent genomic mutations that result from the common oxidatively damaged DNA base, 8-oxo-7,8-dihydro-guanine (OG). MutY enzymes are unusual Base Excision Repair (BER) glycosylases that catalyze the excision of adenine from OG:A bps formed during replication, thereby preventing permanent G:C to T:A transversion mutations. A unique structural hallmark of MutY, along with several other Helix-Hairpin-Helix (HhH) DNA glycosylases, is the presence of a [4Fe-4S] cluster, a cofactor more commonly associated with redox reactions (1–4). Indeed, delineating function in BER for the [4Fe-4S] cofactor has garnered much attention, especially in light of the fact that the glycosylase activity does not require redox chemistry. Moreover, the presence of a redox cofactor within MutY enzymes, whose biological purpose is to prevent oxidative DNA damage-induced mutations, seems too striking to be coincidental.

A wide array of experimental approaches have been used by our laboratory and others to unveil the functional aspects of the [4Fe-4S] cluster in MutY and its human homolog MUTYH (5–12). The cofactor in MutY, and its related HhH DNA glycosylases, is coordinated by four cysteine ligands located within the catalytic domain (Cys-X_6_-Cys-X_2_-Cys-X_5_-Cys, where X is a variable amino acid). The cluster coordination motif sequence and spacing exhibits a high degree of conservation across phylogeny, suggesting that preservation of the cofactor provides crucial functions in these enzymes (13). Structural studies of MutY and Endonuclease III (EndoIII) revealed a DNA recognition role for the [4Fe-4S] cluster via a solvent exposed Iron-Sulfur Cluster Loop (or FCL) motif, corresponding to the “Cys-X_6_-Cys” region, that positions positively charged Lys and Arg side chains for electrostatic interactions with the DNA backbone (6,14). We have also shown that the integrity of the [4Fe-4S] cluster is critical to maintain proper lesion DNA engagement requisite for *in vitro* adenine excision activity (5). However, the absence of the cofactor does not confer changes to secondary structure or thermal stability (6). In *E. coli* MutY, replacements of the cysteine ligands resulted in position- and substitution-dependent impacts on MutY-mediated mutation suppression. For instance, mutation suppression activity was preserved when the second cysteine ligand of the FCL in EcMutY was replaced with Ser, Ala or His, while any kind of substitution of the other Cys ligands compromised this activity (8,10). The [4Fe-4S]^2+^ cluster in MutY, EndoIII, and UDG IV has also been shown to exhibit DNA-mediated redox activity with redox potentials similar to high potential Fe-S proteins (HiPIPs) (7,15). The oxidized [4Fe-4S]^3+^ form of EndoIII was shown to have an increased DNA affinity (7,16,17). These features along with DNA-mediated transport (CT) have been proposed to facilitate DNA lesion localization (18,19). These studies highlight a variety of roles played by the [4Fe-4S]^2+^ cluster cofactor in DNA repair.

We recently revealed an allosteric role for the [4Fe-4S] cluster in MutY enzymes (11). Mapping the locations of cancer-associated variants onto the structure of MUTYH bound to a transition state analog (TSA) containing DNA duplex revealed a hydrogen-bonding network that connects a [4Fe-4S] cluster cysteine ligand, via Arg and Asn residues, to the catalytic Asp. Cancer-associated variants at the Arg-Asn bridging residues disrupt this coordinated network resulting in complete loss of *in vitro* MUTYH glycosylase activity without compromising DNA binding. Indeed, the impact of loss of the network on the activity, structure and dynamics suggests cross-talk between the [4Fe-4S] DNA binding region and the active site. Of note, the structural bridge between the active site and the [4Fe-4S] cluster is also conserved in the HhH BER glycosylases EndoIII and MIG (11). The alterations induced at the active site via an allosteric network provides a compelling explanation for the mechanism by which the [4Fe-4S] redox state of the cofactor may modulate glycosylase activity. Moreover, the link between mutations near the [4Fe-4S] cluster and carcinogenesis highlights the biological significance of the cofactor in the human MUTYH enzyme.

The studies on MutY and MUTYH underscore the importance of its [4Fe-4S] cluster cofactor in its DNA repair function. Nonetheless, mutation-based selection algorithms of evolution not only define rules for structure-activity relationships in biochemical systems, but also dictate exceptions. As such, there are alternative evolutionary variants of EndoIII and MutY lacking the [4Fe-4S]^2+^ cluster. Samrakandi and Pasta (2000) reported the presence of an unconventional MutY in *Streptococcus pneumoniae* which, based on sequence alignments, lacks the [4Fe-4S]^2+^ cluster cysteine ligands (20). In 2002, we also reported variations on the conservation of such ligands in other orthologs of MutY and EndoIII (10). Since then, 45 clusterless MutYs have been identified and phylogenetically characterized, suggesting that clusterless MutY are distributed in two clades; Lactobacillales and variable anaerobic clades (21,22) (**Figure 1D**). The high level of conservation of the [4Fe-4S] cluster in canonical MutYs and phylogenetic clustering of clusterless MutYs indicate a loss of the cofactor in two independent evolutionary events. Representative recombinant clusterless MutYs from *Entamoeba histolytica* and *Lactobacillus brevis* exhibited adenine glycosylase activity and ability to suppress mutations in *E. coli* similar to that of canonical MutYs. Homology structural models of the clusterless MutYs suggested that the absence of the cofactor is compensated for by packing of bulky residues within the cavity that would hold the [4Fe-4S] cluster cofactor in canonical MutYs (21). These results suggest that despite the absence of the [4Fe-4S] cluster, the overall structural scaffold and lesion recognition and excision motifs are conserved in clusterless MutYs.

**Figure 1.**
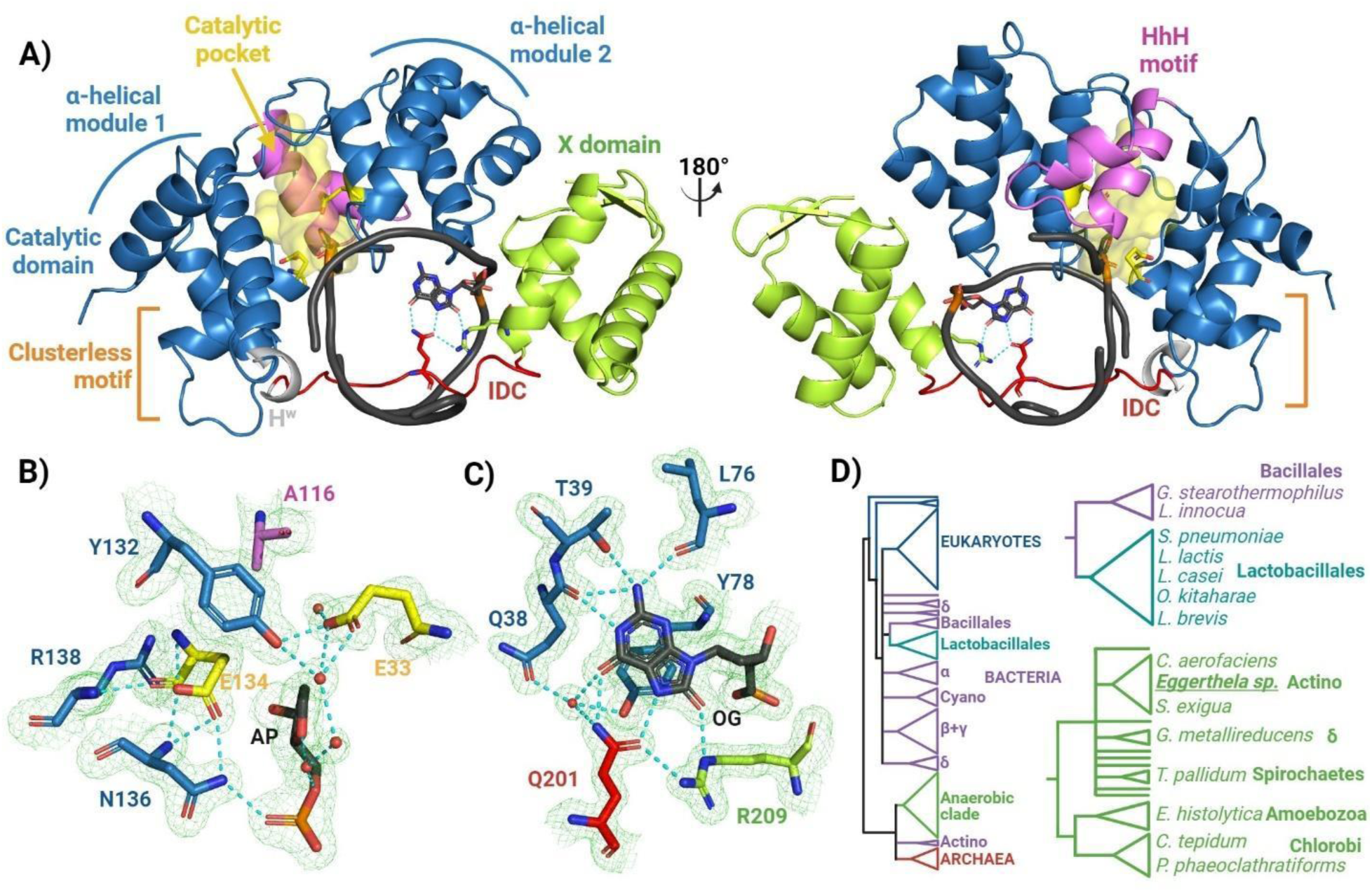
Overview of *Eggerthella sp.* MutYX structure. **A)** MutYX structure in complex with DNA. Each domain and motif are colored differently and indicated with labels. In orange is the region for the [4Fe-4S] cluster in canonical MutYs. **B)** and **C)** Region of MutYX structure highlighting important active site and OG recognition residues, respectively. The 2|Fo|–|Fc| map (green) was calculated to the 1.55 Å resolution limit and contoured at 1.0 rmsd. H-bonds are indicated with cyan dotted lines and water with red small spheres. **D)** Phylogenetic tree of MutY modified from Trasviña-Arenas *et al*, 2016. The clusterless MutY clades are highlighted in green and cyan. Only representative names for Bacillales, Lactobacillales and Anaerobic clades are included. For the full clade members please review reference (21).

In this work, we report the first crystal structure of a clusterless MutY from *Eggerthella sp.* which, contrary to structural homology models, displays complete re-organization of the [4Fe-4S] cluster motif to accommodate its absence, variation in key catalytic residues, and a unique C-terminal OG recognition domain (**Figure 1**). In addition, OG:A specific recognition in clusterless MutYs is provided in a distinct fashion by two key amino acid contacts. Moreover, mutation of these OG recognition residues enhances activity on G:A mismatches. Therefore, given the structural divergence of this type of clusterless MutY from canonical MutYs, we decided to coin it ‘MutYX’.

## RESULTS

### MutYX structure reveals new and conserved features of lesion recognition

Homology models of clusterless MutYs suggested differences in the C-terminal OG recognition domain prompting us to hypothesize that MutY enzymes lacking the [4Fe-4S] cluster evolved a novel mechanism for OG recognition. To explore this hypothesis, we attempted, unsuccessfully, to crystallize the previously reported *E. histolytica* and *L. brevis* clusterless MutYs (21). We then turned to *in silico* evaluation of the crystallization potential for 33 clusterless MutY homologs using the XtalPred algorithm (23) (**Supplementary Figure S1**). Based on these predictions, we designed and cloned the codon-optimized gene for *E. coli* overexpression of *Eggerthella sp. muty* for crystallization trials. We were able to purify the *Eggerthella sp*. MutY in high yields (≈5 mg/L of culture). Moreover, crystallization of the enzyme with a product analog THF:OG-containing 11 nt duplex, resulted in almost immediate (≈2 h) formation of well-diffracting crystals (1.55 Å resolution limit). Given the structural divergence between MutYX and canonical MutY structures deposited in the PDB database, it was not possible to obtain a molecular replacement solution using templates from the PDB. Therefore, an AlphaFold model was generated using the ColabFold server (24,25) to implement molecular replacement with the AlphaFold MutYX model. The structure was solved to yield R/Rfree values of 0.176, 0.197 and deposited in the Protein Data Bank with the ID 8UUC (**Supplementary Table S1**).

The crystal structure of MutYX in complex with the THF:OG-containing DNA duplex displays a modular organization similar to canonical MutYs. The N-terminal catalytic domain and the C-terminal OG recognition domain are connected by an interdomain connector (IDC) region traversing the DNA major groove; MutYX encircles the DNA helix and makes extensive contacts with both DNA strands (**Figure 1A**). Similar to other HhH DNA glycosylases, the N-terminal domain of MutYX harbors the catalytic pocket flanked by two canonical α-helical modules and the HhH motif to enable engagement of the DNA phosphodiester backbone. Indeed, the catalytic domain of *Geobacillus stearothermophilus* MutY (GsMutY; PDB ID 6U7T) and MutYX aligns with a RMSD of 0.943 Å (including 148 α-C atoms).

The most remarkable global differences in the structure of MutYX are the absence of the [4Fe-4S] cluster and the alterations to the C-terminal domain (**Figure 2**). Given the sequence and structural differences of the C-terminal domain of MutYX, we decided to name it the ‘X’ domain. In terms of sequence, the X domain encompasses 67 amino acids, while the corresponding OG recognition domain in MutY includes up to 133 amino acids (**Supplementary Figures S2 and S3**). The overall three-dimensional structure of the C-terminal domain is simpler, being comprised of only three large anti-parallel α-helices and three short anti-parallel β-sheets. (**Figure 2A** and **2B**).

**Figure 2.**
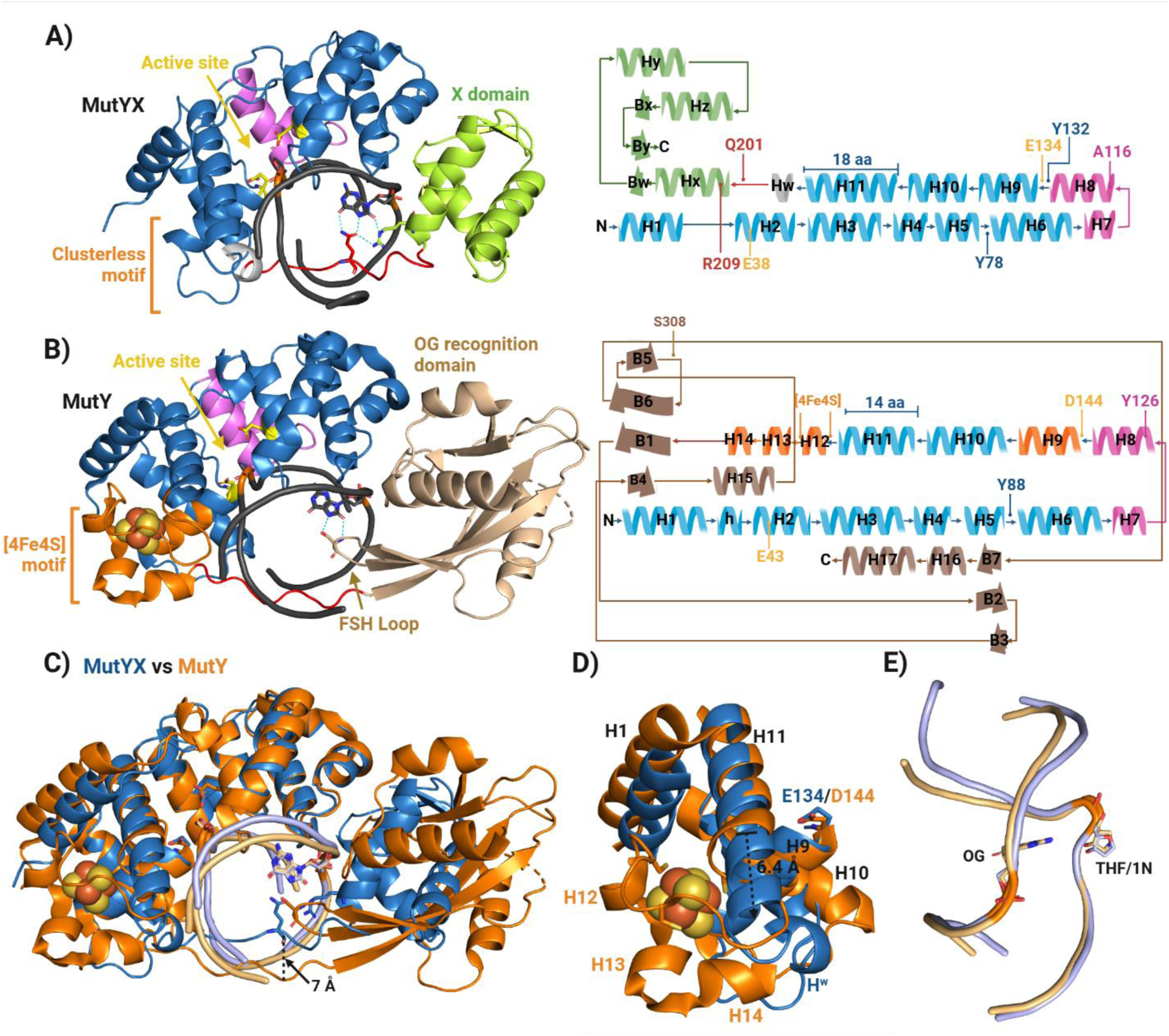
Structural comparison between MutYX and canonical bacterial MutY. **A)** Structural overview of MutYX in complex with THF:OG-containing DNA. On the right panel a map of the secondary structure is displayed highlighting key residues for its activity. **B)** Crystal structure of *G. stearothermophilus* MutY in complex to the transition state analog 1N (PDB 6U7T). On the right panel a map of the secondary structure is displayed highlighting key residues for its activity. **C)** Structural alignment between MutYX (blue) and GsMutY (orange). The internalization of the IDC in MutYX is indicated with a black arrow. **D)** Magnified view of the MutYX vs MutY structural alignment showing the [4Fe-4S] cluster region. The elongation of the α-helix 11 in MutYX s indicated with a dotted arrow. **E)** Comparison of the DNA conformations excreted by their interaction with MutYX (light blue) and MutY (light orange). OG, AP site analogs (THF) and transition state analog (1N) are shown in sticks.

A structural analysis carried out with DALI server (26) found that the X domain shares structural similarities with other subdomains from proteins associated with nucleic acid metabolism, including B-block binding subunit of TFIIIC (RMSD of 6.29 Å) (27), σ appropriation complex (3.91 Å) (28), transcriptional regulator LmrR (5.01 Å) (29) and Cullin (5.01 Å), a scaffold for Ubiquitin ligase (30) (**Supplementary Figure S4**). In contrast, the canonical C-terminal domain has a more complex structure, referred to as the ‘MutT- or NUDT1-like’ domain for its resemblance to the 8-oxodGTPase MutT/NUDT1 (31). In canonical MutYs, the C-terminal domain has a core containing an array of three long anti-parallel β-sheets flanked by α-helices and contains an “FSH” loop that reaches into the DNA major groove to make OG-specific contacts (32). Notably, the X domain in MutYX does not conserve the FSH loop; rather, OG specific recognition is provided by the side chains of Arg209 and Gln201, located within α-helix Hx and the IDC, respectively (**Figure 1C**). Delineation of the additional features of this unusual OG recognition motif are provided in following sections.

Surprisingly, despite the lack of the [4Fe-4S] cluster and changes to the C-terminal domain (**Figures 2C and 2D**), the DNA structure in the complex with MutYX is remarkably similar to that in the *Gs*MutY-DNA complex (**Figure 2E**). The apurinic/apyrimidinic (AP) site analog THF is extrahelically-positioned (**Figures 3A and 3B**), recapitulating the classic nucleotide flipping mechanism of nucleic acid modifying enzymes (31). Tyr78 of MutYX located between the α-helix H5 and α-helix H6 intercalates into the DNA 5’ to the OG, in a mode similar to that observed with Tyr88 in GsMutY (**Figure 3C**). The OG nucleotide within the MutYX-DNA complex structure adopts an *anti* conformation, indicating similar OG:A bp disruption and remodeling following recognition as in GsMutY. The OG base is verified through contacts to both its Watson-Crick and Hoogsteen faces in MutYX and GsMutY (**Figure 3C**). Interestingly, the IDC of MutYX is more closely associated with the DNA than its *Gs*MutY counterpart allowing for its more direct participation in OG recognition (**Figure 2C**). The region surrounding the Gln201 residue is the most closely localized near the DNA, which shows a movement of 7 Å toward OG, relative to the corresponding position in canonical MutYs.

**Figure 3.**
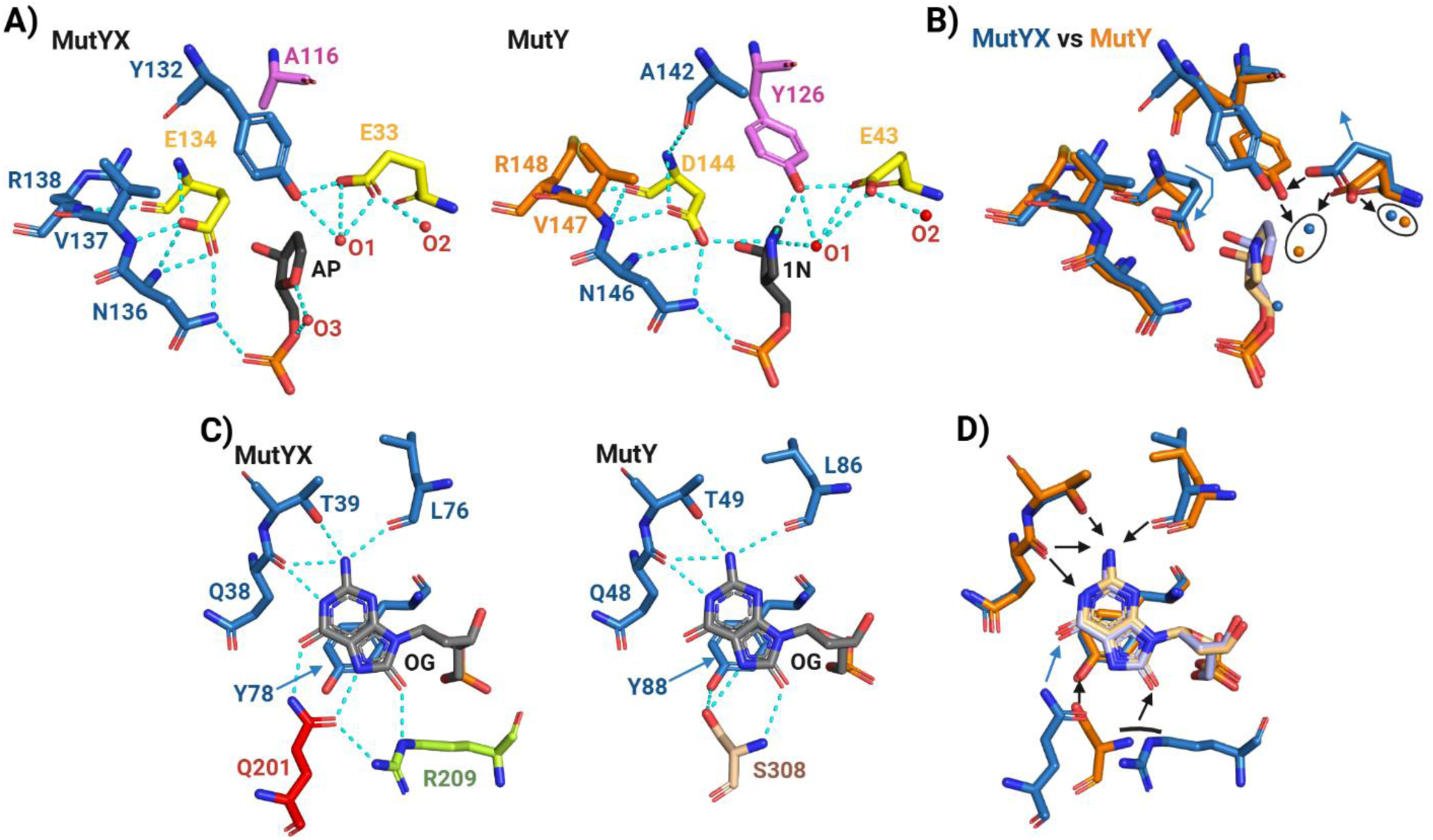
Representation of residues within the active site and the OG recognition sphere. **A)** Configuration of important residues within the active site of MutYX and canonical MutY (GsMutY; PDB 6U7T). Catalytic residues are colored in yellow, residue from the catalytic domain, pink from the Helix-hairpin-Helix (HhH) motif. Water molecules are shown in red small spheres, AP site analogs in black sticks and H-bonds in cyan dotted lines. **B)** Structural alignment of residues from MutYX’s and MutY’s active sites. Conserved water molecules within the active site are circled. Conserved H-bonds are indicated with black arrows and structural displacements or bending are shown with blue arrows. **C)** Residues involved in the OG recognition in MutYX and MutY. The residues from the catalytic domain that interact the Watson-crick face and Hoogsteen face of OG (gray) are in blue sticks. Q201 and R209 residues in MutYX which recognize exclusively the Hoogsteen face of OG from the IDC region and X domain are in red and green, respectively. The canonical S308 from the FSH loop in canonical MutYs are displayed in light brown. **D)** Structural alignment of residues of the OG recognition sphere from MutYX and MutY. Conserved H-bonds are indicated with black arrows while exclusive H-bonds of MutYX are blue arrows.

### Distinctive residues in the catalytic pocket

The high conservation of important residues within the catalytic pocket is a remarkable feature of MutY/MUTYH enzymes (13). Notably, however, MutYX is the first reported MutY homolog with significant variations in active site residues. The most up-to-date catalytic mechanism proposed for MutY requires at least two key catalytic residues, Glu43 and Asp144 (numbering from *Gs*MutY) (12,13,33). The catalytic Glu43 protonates AN7, enhancing its lability, to facilitate N-glycosidic bond cleavage. Asp144 stabilizes the high energy oxocarbenium ion intermediate formed after base excision by formation of a transient covalent acetal intermediate. Finally, Glu43 activates a water molecule for hydrolysis of the acetal intermediate to form the AP site product (13). In MutYX, the catalytic Glu (Glu33) is conserved, however, the catalytic Asp is substituted by another Glu (Glu134) (**Figure 3A** and **3B**). To address the larger size of glutamic acid compared to aspartic acid, the Glu134 side chain of MutYX bends by approximately 136° (considering C^β^ and C^γ^; **Figure S5**) to maintain the same orientation and position of the catalytic Asp144 carboxylate group in GsMutY. Notably, with EcMutY, we have previously observed that WT-like activity in vitro and in cells is preserved if the catalytic Asp is replaced with Glu, illustrating a similar ability to adjust to the longer side chain (34). These results taken together with structural results of MutYX support the notion of mechanism conservation, despite changes in active site residue composition.

The structure of the MutYX active site has many similarities yet also several subtle differences from canonical MutYs. In the MutYX-DNA structure, the AP site analog THF within the active site exhibits a quasi-planar conformation with a slight pucker at C1’ toward C5’ (**Figure S5**). Hence, with precaution due to the resolution limit, we can say that THF adopts a C1’-endo pucker conformation as determined for GsMutY and human MUTYH structures solved with a DNA duplex containing the positively charged (3R,4R)-4-(hydroxymethyl)pyrrolidin-3-ol (1N) nucleotide opposite OG (11,35). Given the lack of positive charge at the C1’ position in THF, it is not surprising that active site residues of MutYX do not make direct contacts to the AP site analog. A water molecule (O3) has the sole interaction with the AP site analog by mediating an H-bonded to the oxygen in THF corresponding to O4’ in an AP site (**Figure 3A**). Water molecules are also observed interacting with the side chains of the catalytic residues Tyr132 and Glu33. Water molecule O1, is interacting with the acidic side chain of Glu33 and the hydroxyl group of Tyr132. A similarly positioned water molecule in the GsMutY structure has been proposed to be the nucleophilic water (35). However, the O1 water in the MutYX structure is farther away from the THF (3.5 Å), likely due to the presence of THF versus 1N (**Figure 3B**). The water molecule O2 is also in close-proximity for H-bonding to Glu33, which also shows a displacement of 1.8 Å relative to the corresponding residue in MutY (Glu43). The triad of Glu43, Tyr126 and the H_2_O molecule O1 that interacts with 1N in GsMutY and their structural conservation in different MutY/MUTYH structures is consistent with their proposed roles during catalysis (11,35,36). Particularly, Tyr126 has been proposed to electrostatically stabilize the charged TS and also position and modulate the acidity of Glu43 (35,37). This Tyr is located at the α-helix H8 and is highly conserved in most MutYs and the HhH BER glycosylases MBD4 and MIG (13). Remarkably, in MutYX, the amino acid at the corresponding position is Ala (Ala116) (**Figure 3A**), however, a different Tyr (Tyr132) is recruited from a helix-helix connector to maintain the H-bond network with the catalytic Glu33 and the water molecule O1, preserving a similar constellation of residues as is in GsMutY. Interestingly, the residue at the sequence position of Tyr132 in MutY is an Ala. This represents an interesting Tyr↔Ala swap in MutY/MutYX homologs to preserve similar catalytic roles for the Tyr residues within the active site.

### MutYX compensates for the absence of the [4Fe-4S] cluster by stabilization of Helix H9

The critical features that are linked to the [4Fe-4S] cluster motif in canonical MutYs suggest that MutYX may have reorganized and acquired new structural elements to preserve enzyme function. The [4Fe-4S] cluster in canonical MutYs is surrounded and shielded from solvent by three α-helices and the FCL motif (**Figure 4**). MutYX does not exhibit structural remnants of the [4Fe-4S] cluster and its FCL motif, except for helix H9 that is positioned close to the [4Fe-4S] cluster in canonical MutYs. The FCL motif in MutYs enables DNA-protein interactions, mainly through electrostatic interactions between its positively charged residues and the phosphodiester backbone (6). For example, in the GsMutY structure, an Arg in the FCL H-bonds directly to the phosphodiester of the third nucleotide downstream (-3) of the TS mimic 1N (**Figure 4A**). In MutYX, a similar interaction is present with Arg193. However, upon loss of the FCL, MutYX acquired a novel α-helical component, Hw, not present in canonical MutYs. The α-helix Hw is positioned near the AP-site analog-containing strand in close proximity to project Arg193 toward the phosphodiester bond at -3 nt to maintain a similar H-bond interaction. Helix Hw is preceded by helix H11 that presents an extension of 4 residues (≈6 Å) in MutYX in comparison to its MutY counterpart (**Figure 2D and 5A**).

**Figure 4.**
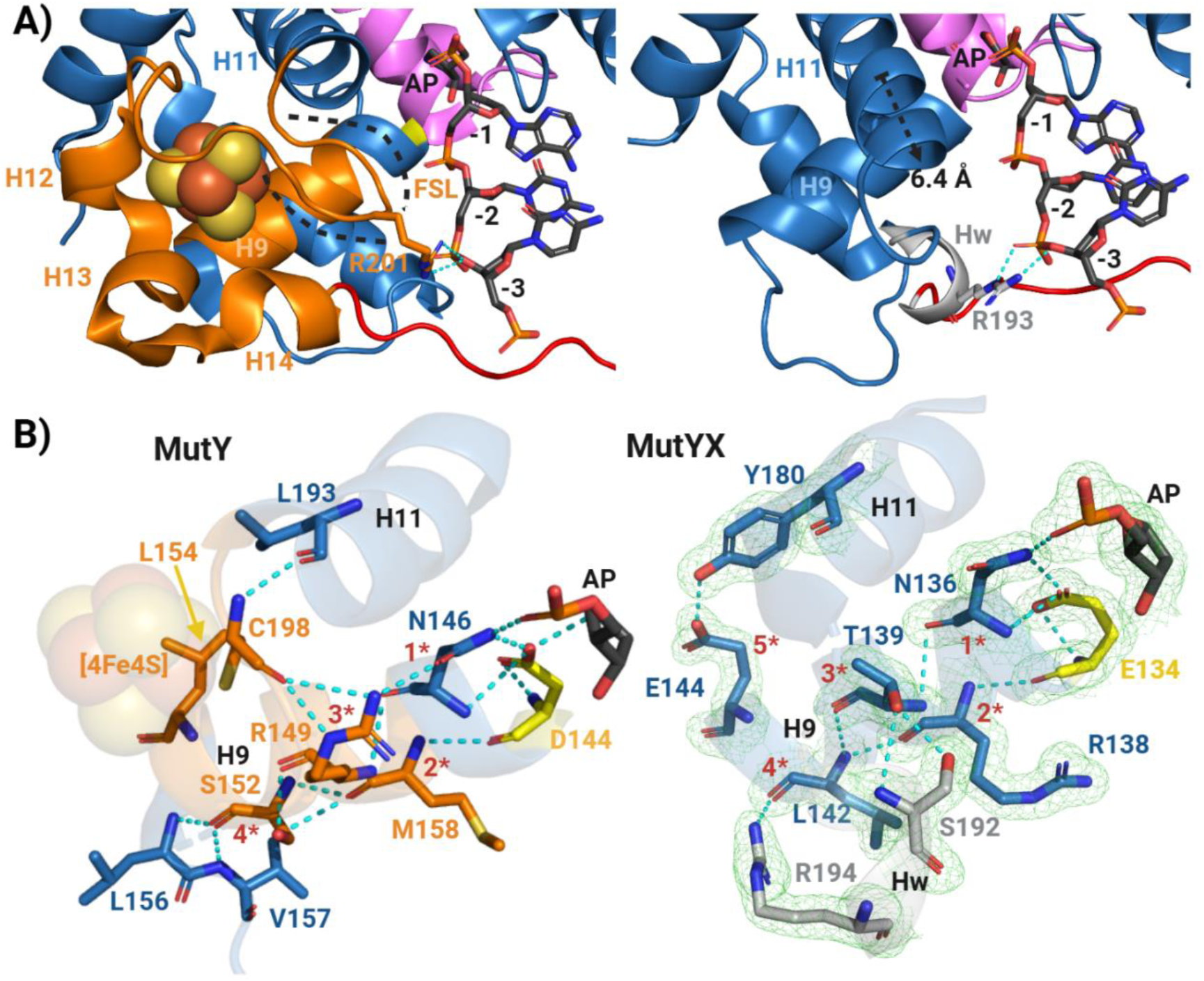
Structural analysis of the [4Fe-4S] cluster motif; its role in DNA-protein interactions and structural dispensability in MutYX. **A)** Close-up of the [4Fe-4S] cluster motif (orange) in MutY (Left panel) and corresponding region of MutYX (Right panel). The [4Fe-4S] cluster is indicated with dotted contour and R201 residue is showcased interacting with the phosphodiester backbone at the third nucleobase (-3) downstream the AP site analog. Similarly, the MutYX’s R193 from the α-helix Hw has the same interaction pattern as MutY’s R201. **B)** MutY α-helix H9 from the [4Fe-4S] cluster motif is stabilized by an intricated H-bond network in MutY (Left panel) mainly through 4 anchoring or stabilization points (red numbers with asterisks) which involves the cysteinyl ligand. In the right panel the reconfiguration of corresponding region of the [4Fe-4S] cluster motif in MutYX is shown. The intricate H-bond network is conserved where the novel α-helix Hw (R194) and the extended H11 (Y180) of MutYX participate as a fourth and fifth anchoring points for α-helix H9 stabilization. The 2|Fo|–|Fc| map (green) was calculated to the 1.55 Å resolution limit and contoured at 1.0 rmsd.

**Figure 5.**
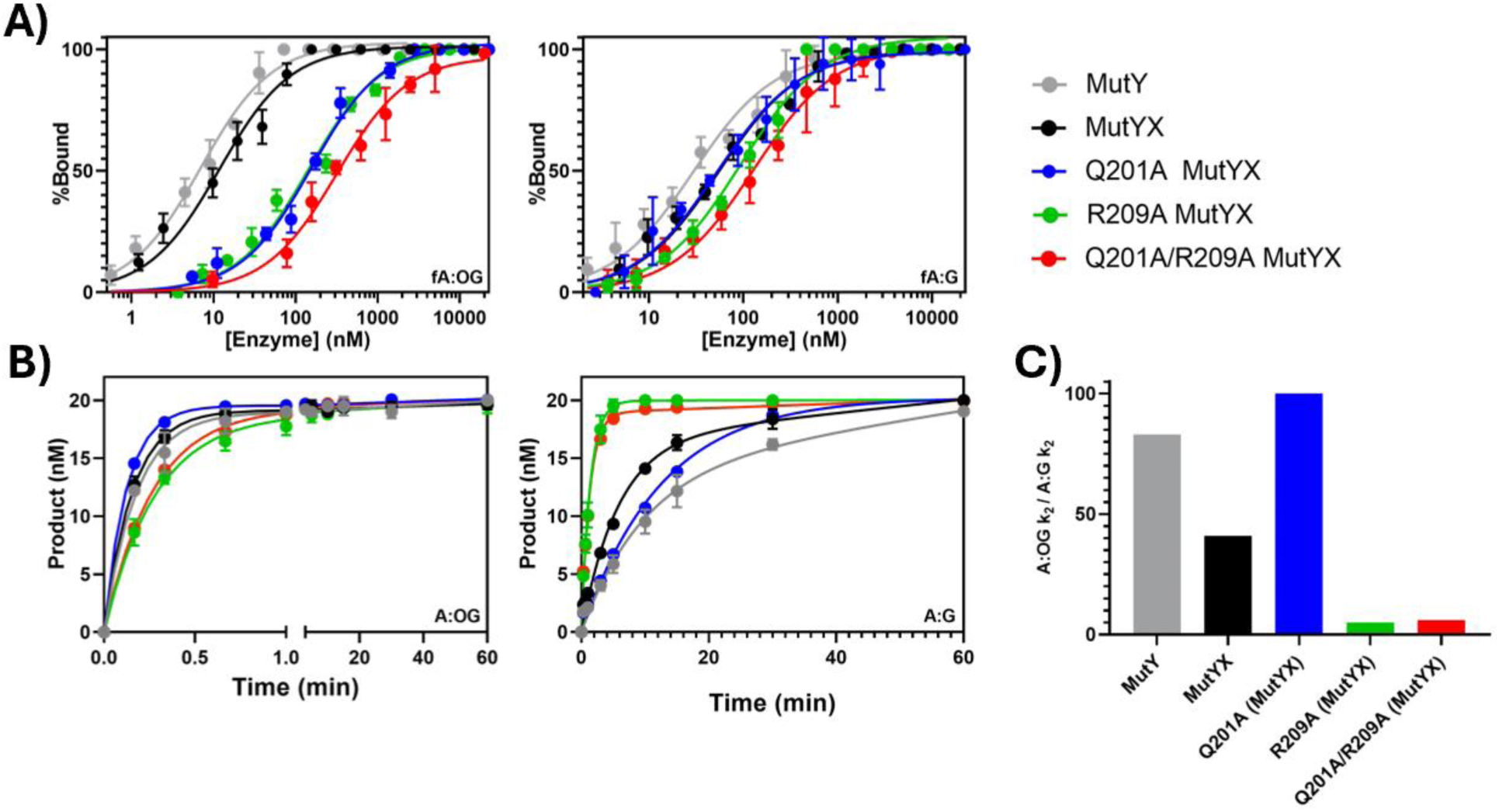
MutYX OG:A lesion affinity and specificity is dictated by Arg209 and Gln 201. **A)** Fluorescence polarization experiments assess relative binding affinities of MutYX and mutants relative to EcMutY for 30-bp DNA duplex containing OG (top left) or G (top right) across from a non-cleavable 2’-fluorine-2’-deoxyadenosine analog (fA). **B)** Single turnover (STO) glycosylase assays determination of adenine excision rate constant (*k_2_*). The STO experiments were performed using 100 nM of active enzyme concentration and 20 nM of DNA substrates at 20 °C. **C)** Plot displaying substrate preference based on the *k_2_*-associated specificity (A:OG *k_2_*/A:G *k_2_*).

We recently revealed an H-bonding network that communicates changes at the DNA binding [4Fe-4S] site to allosterically regulate protonation and positioning of the catalytic Asp in MutY enzymes (11). The network is mediated by two residues Arg149 and Asn146 in helix H9 of GsMutY, that H-bond with each other, and to the cysteine ligand Cys198 and the catalytic Asp144, respectively (**Figure 4B**). Helix H9 is stabilized by 11 H-bonds anchored by residues Asn146, Met158, Arg149 and Ser152. Moreover, the [4Fe-4S] cluster, through its cysteine ligand Cys198, bridges an additional stabilization point involving the Leu193 and Arg149 from the helices H11 and H9, respectively. Therefore, the structural stabilization of H9 is the primary mediator of the functional connectivity. Interestingly, in MutYX, the helix H9 is the sole remaining structural component of the [4Fe-4S] cluster binding motif. As in canonical MutYs, helix H9 is substantially stabilized by 10 H-bond interactions involving 5 residues as anchoring points (**Figure 4B**). In the absence of the [4Fe-4S] cluster, MutYX relies on the helices Hw and H11 which directly H-bond the last portion of the helix H9 (Glu144 and Leu142). Therefore, the extension of helix H11 and acquisition of Hw are the most prominent structural adaptations evolved to stabilize helix H9. Additionally, its structural conservation in canonical and clusterless MutY is evidence that H9 is important to modulate the protonation and positioning of the catalytic carboxylate.

### Gln201 and Arg209 represent a novel OG recognition sphere

OG recognition in MutY homologs is provided by specific H-bond contacts from residues to the Watson-Crick and Hoogsteen faces of OG. In canonical MutYs, the Watson-Crick face of OG is recognized by residues from the catalytic domain (Gln48, Thr49 and Leu86 in GsMutY), while its Hoogsteen face is recognized by a Ser (308 in GsMutY) of the FSH loop within the C-terminal domain (11,32) (**Figure 3C**). In addition, a Tyr residue (88 in GsMutY) intercalates into the DNA helix 5’ to OG to promote adenine extrusion and stabilize the distorted DNA conformation required for catalysis. In MutYX, the Watson-Crick face recognition is conserved and sustained similarly by residues Gln38, Thr39 and Leu76 (**Figure 3C**). Moreover, the positioning of the intercalating Tyr (78 in MutYX) is identical with respect to OG in MutYX. However, the Hoogsteen face of OG is recognized by MutYX in a distinct manner. The side chain of residue Gln201 from the IDC projects towards OG to H-bond with NH7 and O6 of OG. Additionally, the side chain of Arg209 at the end of a helix in the X domain near the IDC is bent with respect to the OG base to allow for NH^ε^ to H-bond directly to the 8-oxo moiety of OG and the NH2^η2^ to H-bonding with the side chain oxygen of Gln201. The coordinated engagement of the Hoogsteen face of OG by Gln201 and Arg209 constitutes a new motif for OG-specific recognition.

### Arg209 provides for unique recognition and high specificity of OG:A

To investigate the unique OG recognition sphere of MutYX, we replaced Gln201 and Arg209 with Ala individually or together for adenine glycosylase and lesion affinity assays. Under multiple turnover conditions (MTO; [E] < [S]), MutYX displayed biphasic kinetic behavior characterized by an initial rapid burst phase of product formation followed by a slower linear phase (**Supplementary Figure S6**). The kinetic behavior is similar to that observed in canonical MutYs as a consequence of the slow release of the AP site DNA product. Accordingly, a similar kinetic scheme and approach as we previously reported was used (38,39) to delineate the kinetic parameters describing glycosidic bond cleavage (*k*_2_) and product release (*k*_3_), and % active fraction (Scheme 1 in methods, **Figure 5A** and **Table 1**). WT MutYX exhibited % active fractions (relative to total protein) that were slightly higher (42%) relative to the Q201A (22%), R209A (30%) and Q201A/R209A (27%), suggesting that perturbations at these positions are functionally impactful. The kinetic parameters related to product release (*k_3_*) determined from the MTO glycosylase assays, indicate similar limited AP site-DNA release after base excision (0.002-0.006 min^-1^; **Figure 5A** and **Table 1**). Initial adenine glycosylase assays were performed with MutYX and EcMutY under single turnover condition (STO; [E]>[S]) using 100 nM active enzyme and 20 nM DNA duplex at 37 °C; however, the reactions were too fast to be measured using manual methods and therefore, the reaction temperature was decreased to 20 °C to obtain measurable and comparable results to those reported previously for other clusterless MutYs (21). At 20 C°, WT MutYX exhibits a similar ability to mediate adenine removal from an OG:A duplex substrate compared to EcMutY, as evidenced by *k*_2_ values of 6.2±0.7 and 5.8±0.2 min^-1^, respectively. R209A and R209A/Q201A MutYX exhibited a decrease in their adenine excision rate constants *k*_2_ (3.4 ± 0.5, 3.6 ± 0.2 min^-1^), consistent with an important role for Arg209 in OG-specific recognition. In contrast, and surprisingly, Q209A MutYX resulted in an *increase* in adenine excision (*k*_2_ = 8.0 ± 0.3 min^-1^); this suggests that removal of the H-bonding interactions mediated by Gln209 may “free up” Arg209 to more easily engage OG to mediate adenine excision.

**Table 1.** Apparent binding (*K_1/2_*), and Glycosylase (*k_2_*) rate constants for *E. coli* MutY and MutYX variants.

| | $K_{1/2}$ (nM)<sup>a</sup> | | $k_2$ (min<sup>-1</sup>)<sup>b</sup> | | | $k_3$ (min<sup>-1</sup>)<sup>c</sup> | |
| --- | --- | --- | --- | --- | --- | --- | --- |
| | OG:fA<sup>FAM</sup> | G:fA<sup>FAM</sup> | OG:A | G:A | $k_2$OG:A/<br>$k_2$G:A | OG:A | G:A |
| EcMutY | 7 ± 1 | 30 ± 3 | 5.8 ± 0.2 | 0.10 ± 0.05 | 83 | 0.001±0.001 | 0.05±0.03 |
| WT MutYX | 12 ± 1 | 52 ± 4 | 6.2 ± 0.7 | 0.15 ± 0.05 | 41 | 0.004±0.001 | 0.003±0.001 |
| Q201A | 156 ± 9 | 53 ± 5 | 8.0 ± 0.3 | 0.10 ± 0.05 | 100 | 0.006±0.001 | 0.003±0.001 |
| R209A | 143 ± 11 | 94 ± 7 | 3.4 ± 0.5 | 0.7 ± 0.1 | 5 | 0.002±0.001 | 0.005±0.001 |
| Q201A/R209A | 315 ± 30 | 129 ± 13 | 3.6 ± 0.2 | 0.7 ± 0.1 | 6 | 0.006±0.001 | 0.006±0.001 |

<sup>a</sup> $K_{1/2}$ determination through fluorescence polarization using 5' 6-FAM labeled DNA substrate containing OG:fA or G:fA lesions performed at 25 °C.
<sup>b</sup> $k_2$ rate constants were determined under single turnover (STO) conditions at 20 °C.
<sup>c</sup> $k_3$ rate constants were determined under multiple turnover (MTO) conditions at 37 °C.

*E. coli* MutY was originally discovered as an adenine glycosylase active on G:A mismatches, and the relevance of this activity is still debated (40,41). Given the alternative OG recognition sphere of MutYX, we were motivated to ascertain the relative substrate specificity (OG:A vs G:A) of WT MutYX. Notably, WT MutYX displays two-fold faster adenine excision with G:A than EcMutY, with *k_2_* values of 0.15 vs 0.07 min^-1^, respectively (**Figure 5B**). Moreover, replacement of Arg209 with Ala (alone or with Gln201) *enhanced* MutYX activity on the G:A substrate. In terms of glycosylase rate-associated specificity (*k_2_* OG:A/ *k_2_* G:A), EcMutY had an 80- fold preference for OG:A substrate over G:A (**Figure 5C**). In contrast, WT MutYX displayed only a 40-fold preference for the OG:A substrate. Most dramatically, the increased activity of the single mutant R209A and double mutant Q201A-R209A of MutYX with G:A resulted in a substantial *decrease* in preference for OG:A over G:A, exhibited by only 5- and 6-fold preference for OG:A, respectively. This suggests that Arg209 drives the specificity of MutYX for OG:A lesions.

The impact of the OG recognition residue mutations of Gln201 and Arg209 in MutYX on lesion recognition was further assessed by measuring the relative dissociation constants (*K_D_)* using electrophoretic mobility shift assays (EMSA) and fluorescence polarization. The relative affinity for the substrate and products was gauged using cleavage-resistant nucleotides, 2’-fluoro-2’-deoxyadenosine (fA) and the abasic site analog, THF, respectively, in 30-bp duplexes positioned opposite OG and G. In the EMSA of WT and Q201A MutYX with the OG:fA substrate analog duplex, we obtained *K_D_*values that are similar to WT EcMutY (*K_D_* of ∼40-100 pM) (**Supplementary Figure S7**). Notably, we were unable to detect significant levels of enzyme-DNA complex with R209A MutYX and the substrate analog duplex, suggesting a significantly reduced affinity and/or fast off-rate. In contrast, using the OG:THF or G:THF duplex, the affinity was high for WT MutYX and all of the variants, such that even at the lowest enzyme concentration the DNA was fully bound (estimated *K_D_*<10 pM). The high affinity for the OG:THF duplex with MutYX is similar to that observed with WT EcMutY; however, EcMutY exhibited reduced affinity for G:THF-containing DNA duplex (260 pM; **Supplementary Figure S7B**). The high affinity for the product analogs is consistent with the slow turnover (*k_3_*) observed with WT, Gln201 and Arg209 MutYX variants with G:A-containing substrate (0.003-0.006 min^-1^; **Table 1, Supplementary Figure S6**), indicating extremely slow product release in all cases with MutYX. With EcMutY, reduced affinity for the G:THF duplex is consistent with higher *k_3_* value (0.05 min^-1^) with G:A substrates (38). Thus, MutYX retains extremely high affinity for AP sites opposite both OG and G. This may be rationalized in part by analysis of the structure that suggests that MutYX makes more H-bonds contacts than GsMutY to the DNA duplex (32 vs 29 respectively, excluding the H-bond contacts with 1N in GsMutY) (**Figure S8**). Interestingly, the increased number of interactions are with the 1N/THF containing strand, rather than the OG strand, resulting in less impact of replacing OG with G.

We further delineated MutYX and EcMutY relative lesion bp affinities utilizing fluorescence polarization assays with the same OG:fA and G:fA-containing 30 bp duplex sequence harboring 6-FAM fluorescent label. Due to detection limits, a higher duplex concentration (500 pM) was needed and resulted in larger apparent *K_D_* (*K_1/2_)* values than those observed in EMSA. However, importantly, the relative *K_1/2_*values highlight the participation of both Arg209 and Gln201 in lesion recognition; single replacements to Ala result in ∼10-fold loss in binding affinity for OG:fA, and double mutation to Ala results in ∼20-fold loss in binding affinity (**Figure 5A** top left). Notably, the binding defects observed with Q201A, R209A, and Q201A/R209A with OG:fA effectively disappear when using the non-preferred substrate (G:fA), such that that are only 1- to 3-fold reduced relative to WT MutYX (**Figure 5A** top right). Altogether, the structure of MutYX and the biochemical data indicate that Arg209 residue dictates MutYX specificity for OG and Gln201 residue plays a subtle role in stabilizing OG in its *anti* conformation, facilitating appropriate positioning of the 8-oxo group for interaction with Arg209. The Q201A mutation may provide more flexibility for the Arg 209 side chain and the OG recognition loop leading to less stringent recognition of the opposition base (OG or G) and potentially aiding in base pair disruption and nucleotide flipping to enhance adenine excision.

## DISCUSSION

The high-resolution (1.55 Å) crystal structure of MutYX in complex with DNA reveals functional and structural insight into the evolutionary divergence from canonical MutYs that contain a [4Fe-4S] cofactor. Indeed, the MutYX structure provides the first experimental structural evidence that illustrates a mechanism of [4Fe-4S] cluster dispensability in BER glycosylases. Previously, homology modeling suggested that clusterless MutYs maintain overall structure and function by stacking of bulky residues within the space occupied by the [4Fe-4S] cluster cofactor in canonical MutYs (21,22). However, the structural reconfiguration to accommodate the absence of the [4Fe-4S] cluster in clusterless MutYX is both complex and subtle, while effectively succeeding to retain OG:A lesion recognition and proper positioning of catalytic residues to preserve base excision repair function. An additional surprise in MutYX is the presence of a unique C-terminal OG recognition domain, that we refer to as the “X” domain to emphasize its distinct structure and OG recognition mode. OG recognition in MutYX relies on two residues, Gln201 and Arg209 that map to the end of the IDC region and the beginning of the X domain, respectively. The OG recognition mode in MutYX is distinct from that of canonical MutYs that use a FSH loop within the C-terminal domain to detect and recognize OG (32). Thus, the MutYX structure also reveals new insights into motifs used for lesion recognition by BER glycosylases.

The OG recognition mode of MutYX manifests in a reduced *in vitro* specificity for OG:A over G:A substrates (38). Remarkably in MutYX OG specificity is dictated primarily by Arg209 which makes H-bond contacts with the 8-oxo moiety of OG. Mutation of the Arg to Ala *enhanced* processing of G:A-containing substrate DNA. Although, MutY is able to process both OG:A and G:A mismatches *in vitro* (38,42), there is no compelling evidence that MutY-mediated G:A repair occurs in a cellular context in *E. coli* (40) or human cells (41). Indeed, in bacteria and eukaryotes avoidance of G:A repair would be expected to be advantageous since this activity would be pro-mutagenic activity in the absence of mechanism to differentiate the daughter versus parental strands during replication. Bacterial mismatch repair (MMR) relies on the identification of hemimethylated DNA to identify the parent strand (43). In human cells, MMR proteins may participate through an OG:A-specific MUTYH activity stimulated by MutSα (44). In a genome-wide search we could not identify MMR recognition and excision proteins MutS, MutH and MutL in genomes of *Eggerthella* genre. We only were able to identify proteins that participate in downstream steps of MMR such as, RecJ, UvrD, DNA polymerase III and SSB (**Table S2**) (45). Although, the MMR pathway seems incomplete, a functional MMR DNA repair pathway may be present in *Eggerthella* with highly divergent homologs of MutS, MutH and MutL that were not detected in our search. Little is known about *Eggerthella* DNA repair mechanisms, however, the fact that it contains MutYX, preserving an OG:A specific adenine glycosylase activity, suggests that other novel repair proteins may be present in these organisms.

In MutY enzymes, the [4Fe-4S] cluster participates in a variety of ways in enzyme function and its presence is required for adenine excision activity (5,8,10). The structure of MutYX reveals structural changes that ensued to compensate for the lack of the [4Fe-4S] cluster in MutYX to preserve OG:A specific detection and adenine excision. The most conspicuous structural adaptations in MutYX are the extension of helix H11, and the acquisition of helix Hw that both flank helix H9 (**Figure 4**). The flanking of helix H9 by H11 and Hw enable the formation of a H-bond network among several residues from these α-helices that function as anchoring points to stabilize helix H9. Therefore, upon [4Fe-4S] cluster loss, such stabilization mechanism of helix H9 allowed the proper positioning of the catalytic Glu required for nucleophilic attack and stabilization of the oxocarbenium intermediate as the covalent acetal during catalysis (13,33). The H-bond network in MutYX is analogous to the H-bonding network that connects the [4Fe-4S] cluster and the base excision active site pocket to regulate the catalytic residue Asp in canonical MutYs (11,33). In addition, the α-helical component Hw in MutYX substitutes for the interactions with DNA mediated by the FCL in MutYs (6), through Arg192, which forms electrostatic interactions with the phosphodiester backbone of the third nucleobase downstream the AP site, as Arg201 from the FCL does in canonical MutY. These structural adjustments in MutYX allow for maintenance of glycosylase activity despite the evolutionary challenge of loss of the [4Fe-4S] cluster.

A captivating feature of canonical MutY and EndoIII glycosylases is the DNA dependent redox activity of the [4Fe-4S] cluster cofactor (9,15,17). The DNA-dependent redox activity of the [4Fe-4S] cluster in MutY/EndoIII has been proposed to enhance DNA lesion recognition (7,9). In this mechanism, cluster redox status modulates DNA affinity and scanning speed, and also provides a means for communication between cluster-containing repair enzymes via DNA mediated charge-transport (CT). However, the inability of MutYX to use redox sensing for DNA lesion recognition suggests that such a mechanism for enhanced and/or coordinated lesion detection is not essential for genome stability in *Eggerthella*; or at least that MutYX has developed alternative strategies to circumvent the lack of redox-mediated CT mechanism. MutYX may rely solely on conventional sliding-hopping mechanisms to locate rare OG:A lesions, similar to that used by other DNA glycosylases that lack a redox cofactor, such as, OGG1 (46,47), UDG (48), NEIL (49), AAG (50), TDG (51), Fpg (52). Notably, *Eggerthella sp.* is an anaerobic gram-negative Bacillus, and therefore, would experience reduced exposure to oxygen radicals. As a consequence, reduced levels of DNA damage may have provided a relaxed evolutionary pressure that led to the structural innovations in MutYX. It may have been evolutionarily advantageous to dispense of the cofactor to reduce the energetic demand to assemble and incorporate Iron-Sulfur clusters into proteins. Additionally, the potential liability of a fragile redox factor may have outweighed its potential benefits; indeed, we have found the [4Fe-4S] cluster in MUTYH can be its “Achilles Heel” providing a locus for many deleterious cancer-associated variants (11). Moreover, degradation of the cluster may lead to release of Fe resulting in DNA damaging Fenton chemistry. The cost versus benefit analysis may have been the tipping point leading to [4Fe-4S] cluster loss during evolution.

Currently, 46 clusterless MutY homologs have been identified (21). Clusterless MutYs are exclusively clustered in two phylogenetic clades (**Figure 1D** and **6**) suggesting that the loss of the [4Fe-4S] cluster has occurred independently twice during MutY evolution, restricted to *Lactobacillales* and a mixed clade which includes some members of anaerobic/microaerophilic actinobacteria, δ-bacteria, spirochaetes phyla and the protist Entamoeba. The structural details of MutYX, the phylogenetic distribution of clusterless MutYs, and MSA allows us to draw additional insights into clusterless MutY evolution (**Figure 6 and Figure S3**). The phylogenetic separation of the Lactobacillales and the mixed anaerobic/microaerophilic clades of clusterless MutYs permits the classification into two structurally distinct MutY adenine glycosylases lacking the [4Fe-4S] cluster cofactors. Lactobacillales MutYs conserves the conventional OG-recognition domain including the FSH loop, therefore, we refer to this group as Clusterless MutYs. However, the MutY-like adenine glycosylases from the anaerobic/microaerophilic clade are more similar to MutYX, harboring the unique OG recognition sphere (Gln201 and Arg209) and sequence conservation of the X domain. Notably, within MutYXs there are interesting differences. For instance, Entamoeba and Chlorobi MutYXs preserved the catalytic Aspartic acid while the Actinobacteria MutYXs, like *Eggerthella* MutYX, employ a Glutamic acid. Moreover, the tyrosine which participates as a stabilization element of the active site through its interaction with the Glutamic acid is mapped in the α-helix H8 in Entamoeba and Clorobi clades. Therefore, the A→Y swapping between α-helix H8 and the helix-helix connector region of the catalytic pocket in *Eggerthella* MutYX is exclusive in actinobacteria clade.

**Figure 6.**
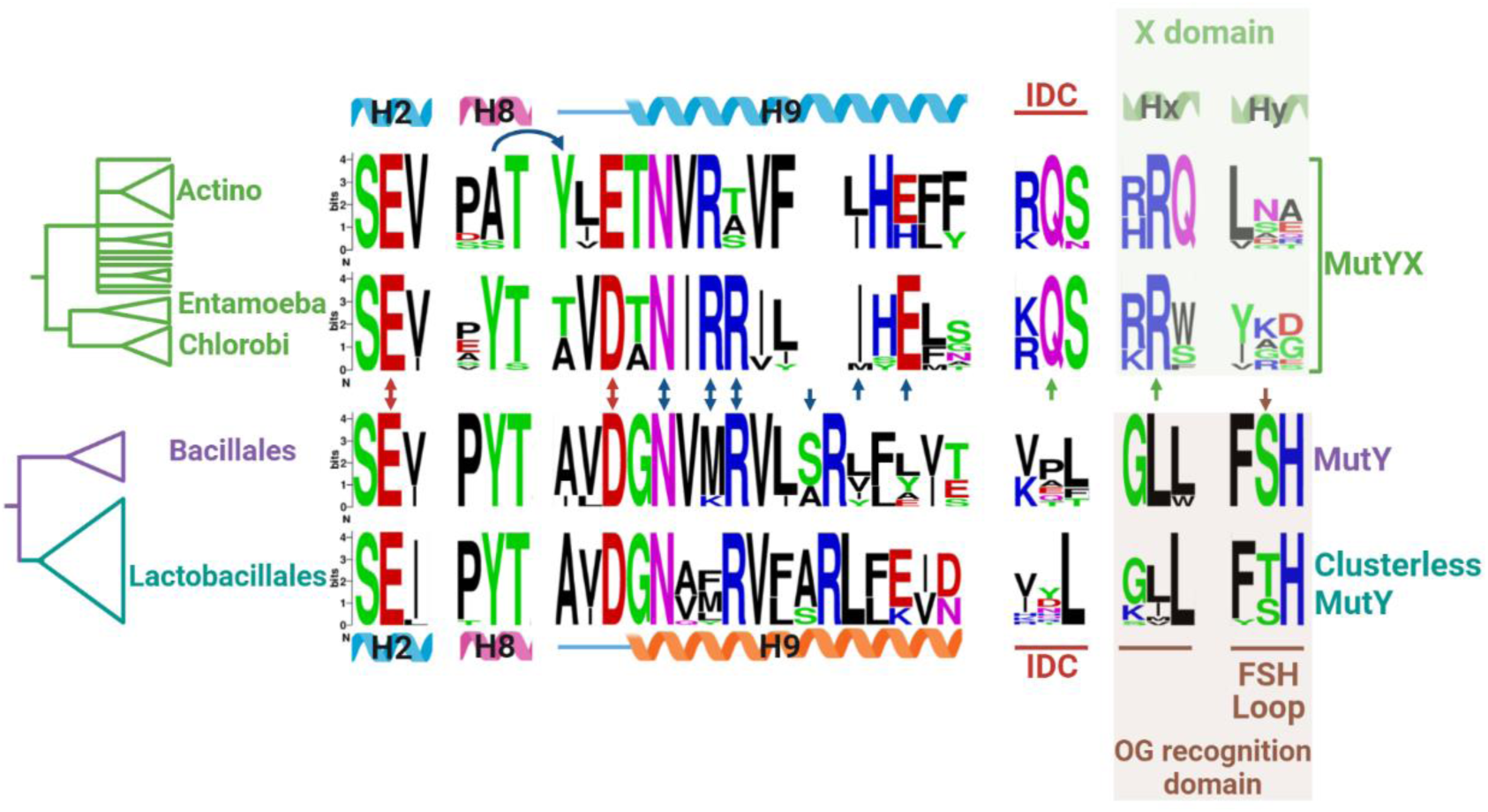
Logo sequence showing the conservation of important residues involved in catalysis and OG recognition in canonical MutY (Bacillale clade), clusterless MutY (Lactobacillus clade) and MutYX (Actinobacteria and Entamoeba/Chlorobi clade). The catalytic residues are indicated with red arrows, α-helix H9 stabilization residues with blue arrows, MutYX’s and MutY’s OG recognition residues with green and brown arrows, respectively. Double headed arrows indicate conservation on both MutYX and canonical MutY, and the direction of the single-headed arrows indicates exclusive conservation. Curved arrow shows the positions involved in the Tyr↔Ala swapping in the active site.

The overwhelming conservation of the [4Fe-4S] in archaeal, bacterial and eukaryotic MutY and its exclusivity in EndoIII, MutY and MIG within the HhH BER superfamily prompted the proposal that the [4Fe-4S] cluster is a late acquisition in BER glycosylase evolution (13). Hence, MutYX is an example of archaic type of MutY prior to [4Fe-4S] cluster acquisition or it is an exemplary case of devolution, whereby evolutionary pressure drove the loss of the metal cofactor in MutYX without losing DNA repair capability. We propose that the later evolutionary scenario is the most likely since (1) the [4Fe-4S] cluster is an ancestral cofactor in MutY conserved in bacteria, archaea and eukaryotes and (2) the phylogenetic clustering of clusterless MutYs and MutYXs subclades are within larger clades containing canonical MutYs harboring the cofactor. In such a case, although synthetic and natural molecular devolution have been described previously (53–56), the crystal structure of MutYX provides atomic details of devolution. It is difficult to conceive the evolutionary pressures that led to the loss of the [4Fe-4S] cluster in MutYXs or Clusterless MutYs. However, some clues come from the biology of non-metallo MutY-containing organisms. For instance, it is documented that *Lactobacillales* phylum harbors a particular group of organisms able to survive at extremely low Fe concentrations (57) that necessitated a series of evolutionary adaptations. One of which may have been dispensing of the use of Fe as a cofactor (58). Similarly, the use as Fe and S may have been evolutionarily exploited in Clorobi phylum and Geobacter, since they harbor Sulfur- and Iron-reducing metabolisms (59,60), respectively.

The structure of MutYX provides an atomic level glimpse at an evolutionary event that compensated the loss of a previously considered indispensable cofactor in MutY and other DNA glycosylases. The distinct structural determinants in MutYX and the ability to manipulate its OG recognition motif to improve specificity to the opposite base, may have practical applications for use as a mutagen or in DNA sequencing (61,62). In addition, the challenges associated with incorporation of the [4Fe-4S] cofactor, and the cofactor’s fragility will also make MutYX attractive as a reagent in various practical applications. The discovery of MutYX therefore provides important insights into evolution of metallo-DNA repair enzymes and is of practical importance in biotechnology.

## METHODS

### *In silico* prediction of clusterless MutY homolog’s crystallizability

The *in sillico* propensity for crystallization of 33 Clusterless MutY homologs was predicted with the XtalPred server (23). The clusterless MutY’s NCBI ID and the results of from the XtalPred analysis prediction are included in **Figure S1**.

### *MutYX* gene cloning and mutagenesis

A codon-optimized *Eggerthella sp. mutyx* gene for *E. coli* overexpression was designed and purchased from Twist Bioscience (San Francisco, California, USA). The ORF lacks the first eighteen codons that were predicted to produce a nonstructured region that was problematic for protein purification. The *mutyx* synthetic gene was cloned into a modified pET28b vector using NdeI and NcoI restriction sites. The modified version of pET28 allows the overexpression of MBP-MutYX protein with a histidine tag at the N-terminal. An internal TEV protease cleavage site was introduced to remove the His-tags and MBP segments from the MutYX protein. Mutagenesis of the pET28-MBP-MutYX construct was carried out by PCR-driven overlap extension (63).

### MutYX overexpression and purification

For MutYX overexpression, a BL21 strain containing pKJE7 vectors was used. The pKJE7 coexpresses dnaK, dnaJ and grgE chaperones to facilitate protein solubility as previously reported (64–66). The BL21(+pKJE7) was transformed with pET28-MBP-MutYX construct and plate onto Luria Broth plates supplemented with 50 μg/mL of Kanamycin and 34 μg/mL of Chloramphenicol. Colonies obtained from the transformation were used to inoculate 2 L of LB media supplemented with the antibiotics previously mentioned and grown at 37 °C/180 rpm until an OD_600nm_ of 0.6. After reaching the aforementioned OD_600nm_, the culture was cooled down for 1 h at 4 °C. The induction of MBP-MutYX fusion protein was carried out by supplementing the culture with 1 mM IPTG. The overexpression was carried out at 15 °C for 16 h. After the overexpression period the bacteria pellets were obtained by centrifugation (6000 rpm/10 min/4°) and stored at -80 °C until needed.

For MutYX purification, the pellets were thawed and resuspended in Lysis buffer (30 mM Tris [pH 8.0], 1 M NaCl and 10% glycerol) supplemented with 1 mM of phenylmethylsulfonyl fluoride. The cellular lysis was carried out by sonication on ice in 20 s cycles using a Branson Sonifier 250 followed by centrifugation at 12,000 rpm for 50 min at 4 °C. The clarified supernatant was incubated with 1.5 mL of Ni^2+^NTA resin (Qiagen) for 1 h at 4 °C with rotation. The slurry was poured over a PD10 column and allowed to flow through via gravity. The protein-loaded resin was washed with at least 25 mL of Lysis buffer followed by 10 mL of elution buffer (30 mM Tris [pH 8.0], 200 mM NaCl, 10% glycerol, and 500 mM Imidazole). To remove the 6xHis-MBP section of the recombinant protein, TEV protease was added to the elution at a ratio of 1:40 w/w (TEV:MBP-MutYX) and dialyzed overnight at 4 °C against Heparin buffer A (30 mM Tris [pH 8.0], 1 mM EDTA, 1 mM DTT and 10% glycerol) supplemented with 200 mM NaCl. Before loading the protein onto a 5 mL Heparin column (Cytiva), the column was equilibrated with buffer A + 100 mM NaCl and the protein diluted with Buffer A to reduce the NaCl concertation to 100 mM. Then, the diluted nickel elution was loaded onto the Heparin column. The loaded heparin column was washed with 25 mL of buffer A + 100 mM NaCl and the elution was carried out with a linear gradient of NaCl (0.1-1M) over 45 min with a flow of 1.5 mL/min using an AKTA FPLC instrument (GE Healthcare). The fractions containing pure MutYX were analyzed by SDS-PAGE and concentrated down using Amicon ultracentrifugation filters (10 000 MWCO). Then, the MutYX protein was subjected to Size-exclusion chromatography using Superdex 200 column with Buffer A+200 mM NaCl. The protein concentration of the elution was estimated by measuring the 280 UV absorbance with an extinction coefficient of 41,035 M^-1^ cm^-1^. Finally, the protein was concentrated again and a portion was aliquoted and stored at -80 °C for biochemical assays and the rest was used for crystallography experiments.

### Preparation of oligonucleotide substrates

The Tetrahydrofuran (THF), OG, G and A containing oligos for crystallography, binding and kinetic experiments were synthesized at the University of Utah core facility. The OG-containing was cleaved from the column and deprotected as previously reported (11). All the oligonucleotides were HPLC-purified, desalted with Sep-Pak C18 desalting cartridge (Waters) and confirmed by matrix-assisted laser-desorption/ionization (MALDI) mass spectrometry at the UC Davis Mass Spectrometry Facility.

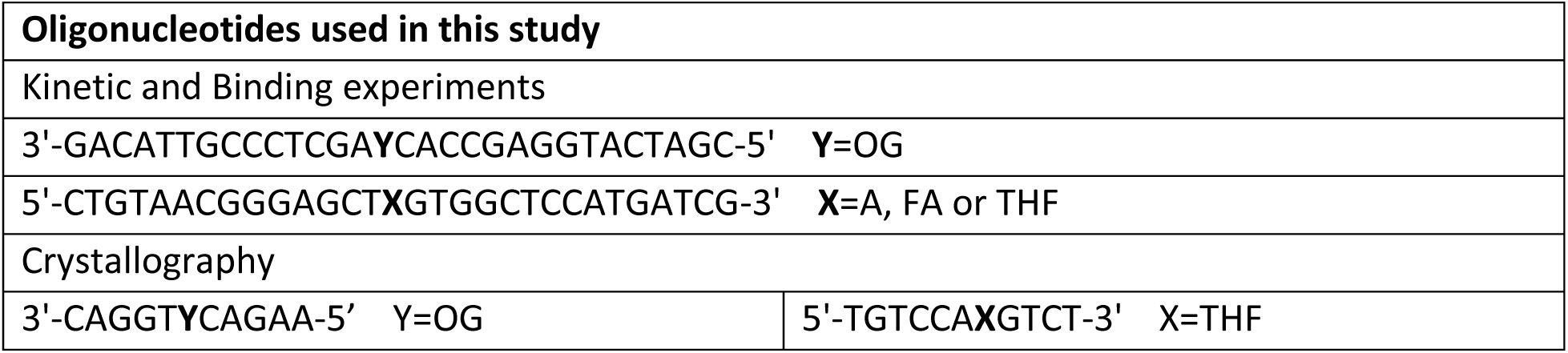

### MutYX crystallization

The MutYX-DNA complex formation for crystallization was carried out with 155 μM and 210 μM of MutYX and THF:OG duplex, respectively. The MutYX-DNA complex was incubated for 20 min at room temperature and mixed with crystallization solutions in 1:1 (v/v) ratio. Large rods crystals of MutYX were obtained overnight by the hanging-drop vapor-diffusion method in 100 mM CAPS/Sodium hydroxide [pH 10.5], 1200 mM Sodium phosphate monobasic/800 mM Potassium phosphate dibasic and 200 mM Lithium sulfate. Before flash cooling, the crystals were transfer to fresh mother liquor supplemented with 30% Ethylene glycol as a cryoprotectant and store in liquid nitrogen. The X-ray diffraction data were collected with 0.2° oscillation on beamline 24-ID-E at the Advanced Photon Source (Argonne National Laboratory).

### Diffraction data processing

Diffraction data from the MutYX-DNA crystals were processed with XDS (67) and scaled with AIMLESS (68). The structure was determined by molecular replacement in PHENIX (69). Given the structural divergence between MutYX and canonical MutY structures deposited in the PDB database, it was not possible to obtain a molecular replacement solution using templates from the PDB. Therefore, an AlphaFold model was generated using the ColabFold server (24,25) resulting in a successful molecular replacement solution. The structure was refined with PHENIX including THF and OG coordination restraints (70). The statistics in data processing and model refinement are shown in supplementary Table S1. The asymmetric unit includes one MutYX molecule complexed with a DNA duplex. All figures depicting structure were generated using PyMOL (The PyMOL Molecular Graphics System, Version 2.0 Schrödinger, LLC.) and Coot (71).

### Glycosylase assay and binding experiments

The glycosylase (*k_2_*) and turnover (*k_3_*) rate constants of MutYX with the OG:A and G:A 30 bp substrates were measured under single turnover and multiple turnover conditions (STO and MTO), respectively as previously reported (39,72), using a minimal kinetic scheme as described below.

**Scheme 1:**
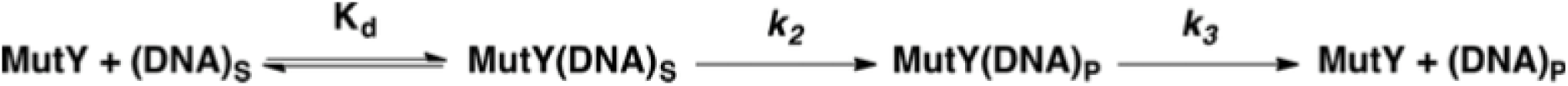

Rate constants (*k_2_)* were determined under single turnover (STO) conditions employing 100 nM active enzyme and 20 nM OG:A- or G:A-containing DNA substrates at 20 °C. Rate constants (*k_3_*) were determined under multiple turnover (MTO) conditions employing 5-8 nM total enzyme and 20 nM OG:A-containing DNA substrate at 37 °C. In order to observe a significant burst with the G:A substrate, the total enzyme concentration was raised to 100 nM to measure *k*_3_ constants for all MutYX variants.

The binding affinity (*K_D_*) was determined using the Electrophoretic Mobility Shift Assays (EMSA) with a noncleavable 2’-fluoro-2’deoxyadenosine (fA) nucleotide paired OG or G (10 pM final concentration), as previously reported (39). The apparent binding affinities (*K_1/2_*) for all MutYX variants and EcMutY were performed using fluorescence polarization to detect the enzyme-DNA complex. The DNA substrates were the same fA:OG and fA:G containing 30-bp duplexes utilized for EMSA, with the exception of possessing a 5’ 6-FAM label. The fluorescence polarization readout was carried out with 500 pM of fluorescently-labeled DNA duplex titrated with increasing concentrations of EcMutY or MutYX. Enzyme concentrations ranged from 1 nM– 20 uM based on Abs_280nm_ and not normalized for differences in activity to avoid masking OG recognition defects within variants. After enzyme titration, steady state enzyme-DNA complex formation was achieved by incubation for 30 minutes at 25 °C. Subsequently, titrants were transferred to a 384 well plate to assess changes in fluorescence polarization on a BMG Labtech Clariostar multimode plate reader with an excitation/emission wavelength of 482/530 nm and a gain adjustment set to 60. All the experiments were carried out in triplicate. The binding and kinetic constants were determined using the GraphPad Prism 7 software.

### Multiple sequence analysis

Sequence alignments were performed with MUSCLE algorithm (73) in the software Geneious (version 4.8.5) (74). The logo sequence was generated in the WebLogo webpage (75).

## DATA AVAILABILITY

Coordinates and structure factors have been deposited in the Protein Data Bank, www.pdb.org (PDB ID code 8UUC).

## ACKNOWLEDGMENTS

We thank Dr. Martin Horvath and Danielle Yama (U of Utah) for pointing out the similarity of the X domain to sub-domains in other nucleic acid binding proteins.

