## Supplementary Information for "Crystal structure of MutYX: A novel clusterless adenine DNA glycosylase with a distinct C-terminal domain and 8-Oxoguanine recognition sphere"

Sheila S. David. University of California, Davis, One Shields Avenue, Davis, CA 95616, U.S.A.

.

C.H. Trasviña-Arenas. Centro de Investigación Sobre el Envejecimiento, Centro de Investigación y de Estudios Avanzados del Instituto Politécnico Nacional (Cinvestav), Unidad Sede Sur, Calzada de los Tenorios, No. 235, Col. Rinconada de las Hadas, Mexico City CP 14330, Mexico

**TABLE OF CONTENTS**

**Figure S1.** Results obtained from XtalPred analysis to predict the likelihood for crystallization of clusterless MutY and MutYX homologs

**Figure S2.** Sequence and secondary structure representation of MutYX. This scheme was generated in PDBsum

**Figure S3.** Multiple sequence alignment. It includes *Eggerthella sp.* MutYX, *G. stearothermophilus* MutY (GsMutY), *L. brevis* clusterless MutY (LbMutY), *E. coli* MutY (EcMutY) and human MUTYH.

**Figure S4.** Representative results with the structural hits for X domain found with DALI server

**Figure S5.** Structural alignment of the AP site analog (THF) from the MutYX structure and the transition state analog (1N) from GsMutY

**Figure S6.** Multiple turnover (MTO) kinetics for determination of turnover rate and active fraction

**Figure S7.** Electrophoretic mobility Shift Assays (EMSA) with WT and OG-recognition motif variants of MutYX.

**Figure S8.** Scheme of DNA-protein contacts in GsMutY and MutYX structures

**Table S1.** MutYX structure diffraction data and refinement statistics

**Table S2.** Results of the search of MMR components in *Eggerthella* genomes.

| NCBI ID | # | [MutY homolog] | EP-class | RF-class | Length | Gravy index | Instability index [II] | Isoelectric point [pI] | Coiled coils | Longest disorder | Percentage of coil structure | Transmembrane helices [TM] | Signal peptides [SP] | Insertions score | Homologs in NR [clustered to 60% | Homologs in PDB |
|---|---|---|---|---|---|---|---|---|---|---|---|---|---|---|---|---|
| BAK44394.1 | 1 | Eggerthella_sp._YY7918 | 2 | 10 | 291 | -0.22 | 34.87 | 5.56 | 0 | 19 | 37 | No | No | 0.08 | 1155 | 32 |
| EEZ61277.1 | 2 | Slackia_exigua | 3 | 10 | 293 | -0.35 | 46.83 | 5.72 | 0 | 19 | 40 | No | No | 0.01 | 1160 | 28 |
| WP_014455772.1 | 3 | Spirochaeta_africana | 4 | 11 | 276 | -0.35 | 56.63 | 7.96 | 0 | 14 | 35 | No | No | 0.08 | 1155 | 32 |
| WP_013709462.1 | 4 | Coriobacterium_glomerans | 4 | 10 | 299 | -0.35 | 58.98 | 8.36 | 0 | 10 | 34 | No | No | 0.14 | 1181 | 29 |
| WP_013945004.1 | 5 | Treponema_paraluiscuniculi | 4 | 11 | 277 | -0.2 | 57.75 | 8.96 | 0 | 15 | 37 | No | No | 0.06 | 1206 | 32 |
| WP_010881791.1 | 6 | Treponema_pallidum | 4 | 11 | 277 | -0.22 | 57.21 | 8.96 | 0 | 16 | 37 | No | No | 0.09 | 1214 | 32 |
| ABB31885.1 | 7 | Geobacter_metallireducens | 4 | 8 | 285 | -0.24 | 52.04 | 9.23 | 0 | 22 | 39 | No | No | 0.02 | 1202 | 32 |
| WP_013969027.1 | 8 | Treponema_caldaria | 5 | 7 | 279 | -0.47 | 51.93 | 9.32 | 21 | 15 | 36 | No | No | 0.06 | 1169 | 32 |
| WP_010942258.1 | 9 | Geobacter_sulfurreducens | 5 | 11 | 285 | -0.4 | 54.11 | 9.51 | 0 | 22 | 35 | No | No | 0.01 | 1137 | 31 |
| CCJ35653.1 | 10 | Methanoculleus_bourgensis_MS2 | 5 | 11 | 348 | -0.43 | 52.59 | 7.72 | 21 | 57 | 47 | No | No | 0.05 | 1170 | 33 |
| NP_662053.1 | 11 | Chlorobium_tepidum | 3 | 8 | 273 | -0.27 | 51.65 | 6.17 | 0 | 16 | 35 | No | No | 0.13 | 1176 | 32 |
| WP_012508226.1 | 12 | Pelodictyon_phaeoclathratiforme | 3 | 4 | 278 | -0.39 | 56.91 | 6.27 | 0 | 11 | 35 | No | No | 0 | 1185 | 32 |
| YP_001998713 | 13 | Cryptosporidium_parvum | 4 | 8 | 277 | -0.29 | 57.44 | 6.37 | 0 | 17 | 35 | No | No | 0.1 | 1187 | 32 |
| WP_006365894.1 | 14 | Chlorobium_ferrooxidans | 4 | 11 | 272 | -0.48 | 56.38 | 9.01 | 0 | 16 | 34 | No | No | 0.02 | 1208 | 32 |
| WP_015711266.1 | 15 | Treponema_azotonutricium | 5 | 11 | 270 | -0.57 | 58.87 | 9.03 | 0 | 15 | 36 | No | No | 0.07 | 1207 | 32 |
| XP_004260533.1 | 16 | Entamoeba_invadens | 5 | 11 | 318 | -0.46 | 44.37 | 9.66 | 0 | 57 | 42 | No | No | 0.01 | 1189 | 31 |
| XP_001740351.1 | 17 | Entamoeba_dispar | 5 | 11 | 307 | -0.59 | 59.51 | 9.41 | 38 | 46 | 38 | No | No | 0.01 | 1240 | 31 |
| XP_653060.1 | 18 | Entamoeba_histolytica | 5 | 11 | 307 | -0.6 | 59.11 | 9.45 | 38 | 46 | 37 | No | No | 0 | 1238 | 31 |
| XP_008857367.1 | 19 | Entamoeba_nutalli | 5 | 11 | 307 | -0.6 | 59.8 | 9.41 | 38 | 46 | 37 | No | No | 0 | 1236 | 31 |
| WP_010290796.1 | 20 | Leuconostoc_pseudomesenteroides | 2 | 5 | 344 | -0.42 | 48.53 | 6.41 | 0 | 10 | 38 | No | No | 0.01 | 970 | 32 |
| WP_002828104.1 | 21 | Weissella_paramesenteroides | 3 | 8 | 366 | -0.43 | 41.83 | 5.7 | 0 | 11 | 41 | No | No | 0.12 | 973 | 32 |
| WP_007745292.1 | 22 | Oenococcus_kitaharae | 3 | 11 | 373 | -0.36 | 41.14 | 5.23 | 0 | 13 | 41 | No | No | 0.12 | 960 | 32 |
| WP_008856803.1 | 23 | Lactobacillus_kisonensis | 4 | 11 | 372 | -0.47 | 45.97 | 8.67 | 0 | 11 | 39 | No | No | 0.12 | 975 | 32 |
| ARW51535.1 | 24 | Lactobacillus_brievis | 4 | 7 | 379 | -0.55 | 46.05 | 7.76 | 0 | 10 | 39 | No | No | 0.11 | 971 | 50 |
| WP_000886134.1 | 25 | Streptococcus_sanguinis | 4 | 9 | 386 | -0.56 | 46.71 | 5.16 | 0 | 10 | 40 | No | No | 0.13 | 966 | 31 |
| WP_004561528.1 | 26 | Lactobacillus_hilgardii | 4 | 8 | 370 | -0.55 | 45.82 | 8.37 | 0 | 10 | 39 | No | No | 0.11 | 969 | 50 |
| WP_000886142.1 | 27 | Streptococcus_mitis | 4 | 7 | 391 | -0.48 | 44.6 | 5.01 | 0 | 12 | 42 | No | No | 0.12 | 967 | 31 |
| WP_000886146.1 | 28 | Streptococcus_pneumoniae_1 | 4 | 1 | 391 | -0.48 | 47.77 | 4.86 | 0 | 12 | 42 | No | No | 0.13 | 969 | 31 |
| WP_010010046.1 | 29 | Lactobacillus_coryniformis | 1 | 2 | 371 | -0.23 | 37.87 | 6.33 | 0 | 12 | 41 | No | No | 0.11 | 978 | 32 |
| ZP_12649903.1 | 30 | Lactobacillus_casei | 1 | 5 | 367 | -0.21 | 37.77 | 6.51 | 0 | 14 | 40 | No | No | 0.1 | 961 | 34 |
| EFQ57131.1 | 31 | Streptococcus_downei_F0415 | 3 | 6 | 389 | -0.53 | 39.61 | 6.51 | 0 | 21 | 42 | No | No | 0.12 | 960 | 32 |
| WP_003042132.1 | 32 | Streptococcus_anginosus | 4 | 8 | 389 | -0.41 | 46.64 | 4.99 | 0 | 15 | 41 | No | No | 0.13 | 964 | 32 |
| WP_015285466.1 | 33 | Methanoregula_formicicum | 4 | 10 | 288 | -0.31 | 48.33 | 9.25 | 0 | 21 | 40 | No | No | 0.07 | 1165 | 32 |

RF crystallization classes
↓ Most promising

1 2 3 4 5 6 7 8 9 10 11

Least promising ↑

EP crystallization classes
↓ Most promising

1 2 3 4 5

↑ Least promising

**Figure S1.** Results obtained from XtalPred analysis to predict the likelihood for crystallization of clusterless MutY and MutYX homologs

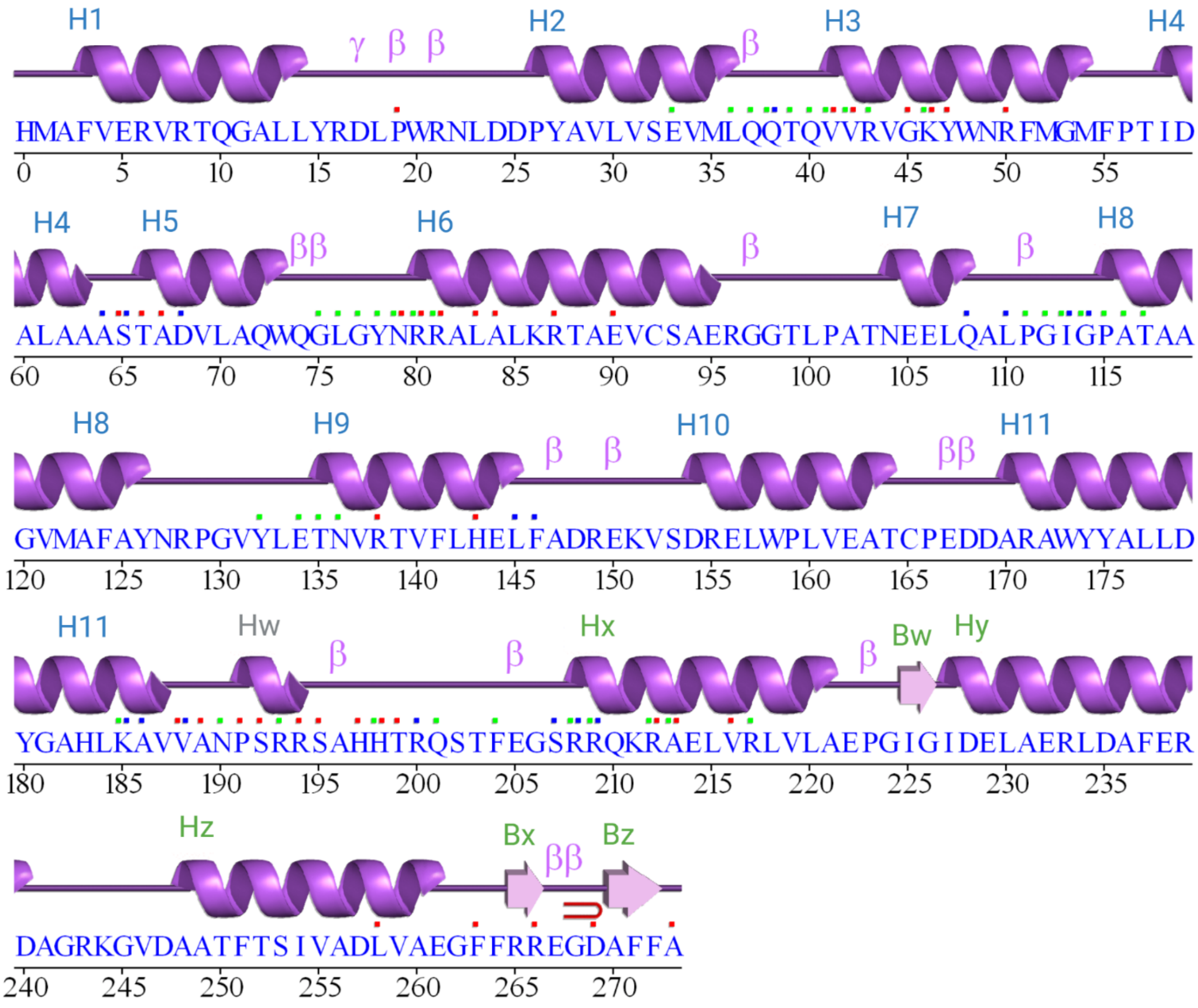

**Figure S2.** Sequence and secondary structure representation of MutYX. This scheme was generated in PDBsum (1). Red, green and blue dots represent residues contact to ligand, DNA and metals, respectively.

```
        1        10        20        30        40        50        60
        |        |         |         |         |         |         |
 MutYX  HMRVTDSEAALWPDGGLSKEAF--------------------------------------
   GsY  MTRETERFP---------------------------------------------------
LbMutY  LIHMVEWTP---------------------------------------------------
EcMutY  ------------------------------------------------------------
 MUTYH  MTPLVSRLSRLWAIMRKPRAAVGSGHRKQAASQEGRQKHAKNNSQAKPSACDACAGMIAE
        61       70        80        90        100       110       120
        |        |         |         |         |         |         |
 MutYX  ------------------------------VERVRTQGALL-------YRDLPWRN----
   GsY  --------------------------------AREFQRDLLDWFAR-ERRDLPWRK----
LbMutY  ------------------------------EKIVAFQETLLKWYDN-NKRNLPWRR----
EcMutY  ------------------------------MQASQFSAQVLDWYDKYGRKTLPWQI----
 MUTYH  CPGAPAGLARQPEEVVLQASVSSYHLFRDVAEVTAFRGSLLSWYDQ-EKRDLPWRRRAED
        121      130       140       160       170       180       190
        |        |         |         |         |         |         |
 MutYX  ----LDDPYAVLVSEVMLQQTQVVRVGKYWNRFMGMFPTIDALAAASTADVLAQWQGLGY
   GsY  ----DRDPYKVWVSEVMLQQTRVETVIPYFEQFIDRFPTLEALADADEDEVLKAWEGLGY
LbMutY  ----DHDPYHIWISEIMLQQTQVQTVIPYYERFMKLFPTVQALASADEAILMKAWEGLGY
EcMutY  ----DKTPYKVWLSEVMLQQTQVATVIPYFERFMARFPTVTDLANAPLDEVLHLWTGLGY
 MUTYH  EMDLDRRAYAVWVSEVMLQQTQVATVINYYTGWMQKWPTLQDLASASLEEVNQLWAGLGY
        191      200       210       220       230       240       250
        |        |         |         |         |         |         |
 MutYX  NRRALALKRTAEVCSAERGGTLPATNEELQA-LPGIGPATAAGVMAFAYNRPGVYLETNV
   GsY  YSRVRNLHAAVKEVKTRYGGKVPDDPDEFSR-LKGVGPYTVGAVLSLAYGVPEPAVDGNV
LbMutY  YSRARNLQKAAQQIVNDYNGQWPTTVKELQE-LSGIGPYTAGAIASIAFNKPVPAVDGNA
EcMutY  YARARNLHKAAQQVATLHGGKFPETFEEVAA-LPGVGRSTAGAILSLSLGKHFPILDGNV
 MUTYH  YSRGRRLQEGARKVVEELGGHMPRTAETLQQLLPGVGRYTAGAIASIAFGQATGVVDGNV
        251      260       270       280       290       300       310
        |        |         |         |         |         |         |
 MutYX  RTVFLHELFADRE---KVSDRELWPLVEATCPEDDARAWYYALLDYGAHLKAVVANPSRR
   GsY  MRVLSRLFLVTDDIAKPSTRKRFEQIVREIMAYENPGAFNEALIELG----ALVCTPRRP
LbMutY  LRVFARLLEIDEDIAKPQTRKLFENIIKKLMPKNRPGDFNQAIMDLG----ASYMSAKNY
EcMutY  KRVLARCYAVSGWPGKKEVENKLWSLSEQVTPAVGVERFNQAMMDLG----AMICTRSKP
 MUTYH  ARVLCRVRAIGADPSSTLVSQQLWGLAQQLVDPARPGDFNQAAMELG----ATVCTPQRP
        311      320       330       340       350       360       370
        |        |         |         |         |         |         |
 MutYX  SAHHTRQSTFEGS-----------------------------------------------
   GsY  SCLLCPVQAYCQA-----------------------------------FAEGVAEE----
LbMutY  DSENSPVKLFNQA-----------------------------------YLDGVEDN----
EcMutY  KCSLCPLQNGCIA-----------------------------------AANNSWAL----
 MUTYH  LCSQCPVESLCRARQRVEQEQLLASGSLSGSPDVEECAPNTGQCHLCLPPSEPWDQTLGV
        371      380       390       400       410       420       430
        |        |         |         |         |         |         |
 MutYX  ----------RRQKRAELVRLVLAEPG---------------------------------
   GsY  --LP-VKMKKTAVKQVPLAVAVLADDE---GRVLIRKRDSTGLLANLWEFPSCETDGADG
LbMutY  --YP-VKTKKKRPIPVNYFGLLIHSQD----DYLFERRPNSGILSRFWMFPLIKGDDIQT
EcMutY  --YPGKKPKQTLPERTGYFLLLQHEDE-----VLLAQRPPSGLWGGLYCFPQFADEES--
 MUTYH  VNFP-RKASRKPPREESSATCVLEQPGALGAQILLVQRPNSGLLAGLWEFPSVTWEPSEQ
        431      440       450       460       470       480       490
        |        |         |         |         |         |         |
 MutYX  ------------------------IGIDE-LAER--LDAFER------------------
   GsY  KEK------LEQMVGE---QYGLQVELTEPIVSF--EHAFSHLVWQLTVFPGRLVHGGPV
LbMutY  KKDASEDDVLRALEAQFLSTYQLEIHVKK-IGGRPVTHTFTHQKWQITLLEAELNNSSDL
EcMutY  ---------LRQWLAQ---RQIAADNLTQ-LTAF--RHTFSHFHL---------------
 MUTYH  LQRKALLQELQRWAG-----PLPATHLRH-LGEV--VHTFSHIKLTYQVYGLALEGQTPV
        491      500       510       520       530       540       550
        |        |         |         |         |         |         |
 MutYX  ---DAGRKGVDAATFTSIVADLVAEGFFRREGDAFFA-----------------------
   GsY  EE-P--YRLAPEDELKAYAFPVSHQRVWREYKEWASG-----------------------
LbMutY  SYFP--GKWISESDFREIAFTKVQTKMWERYQQQKEQ------*LELA*PLG--------
EcMutY  DIVP---MWLPVSSF-TGCMDEGNALWYNLAQPPSVG---------LAAPVERL------
 MUTYH  TTVPPGARWLTQEEFHTAAVSTAMKKVFRVYQGQQPGTCMGSKRSQVSSPCSRKKPRMGQ
```

```
         551       560        570
         |         |          |
 MutYX   ------------------------
   GsY   -------------------VRPPP
LbMutY   --------------ASKRVLRGFL
EcMutY   --------------LQQLRTGAPV
 MUTYH   QVLDNFFRSHISTDAHSLNSAAQ*
```

**Figure S3** Multiple sequence alignment. Sequences include those of *Eggerthella sp.* MutYX, *G. stearothermophilus* MutY (GsMutY), *L. brevis* clusterless MutY (LbMutY), *E. coli* MutY (EcMutY) and human MUTYH. Important residues which are discussed in this paper are colored as follows. Catalytic residues, red; Stabilization Tyr, blue; MutYX's OG recognition, green; MutY/MUTYH's OG recognition, brown; Cysteine ligands for [4Fe-4S] cluster coordination, orange; Arg involved in DNA-protein interactions, purple.

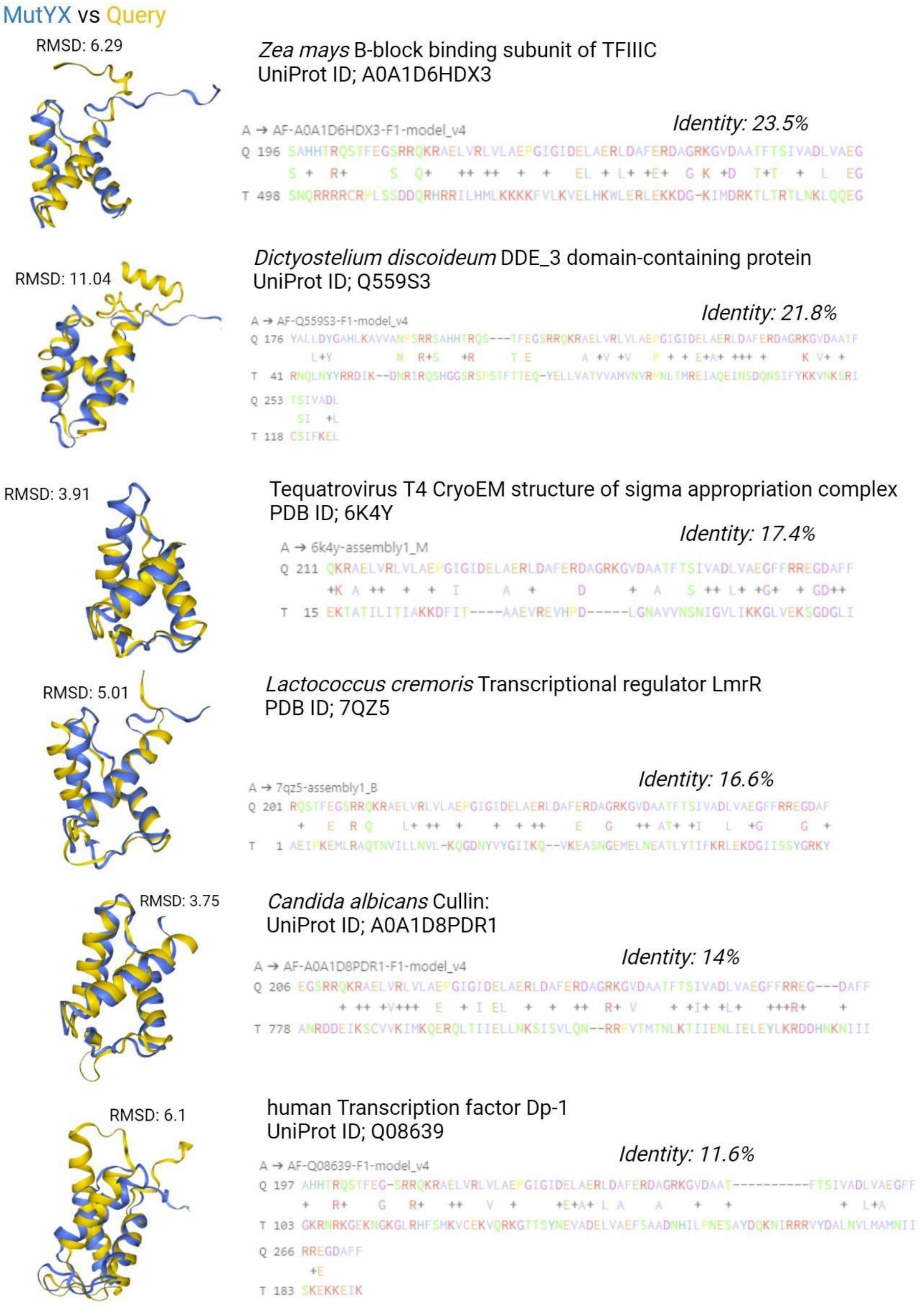

**Figure S4.** Representative results with the structural hits for X domain found with DALI server (2). These representative structural elements were selected based on its amino acid sequence identity.

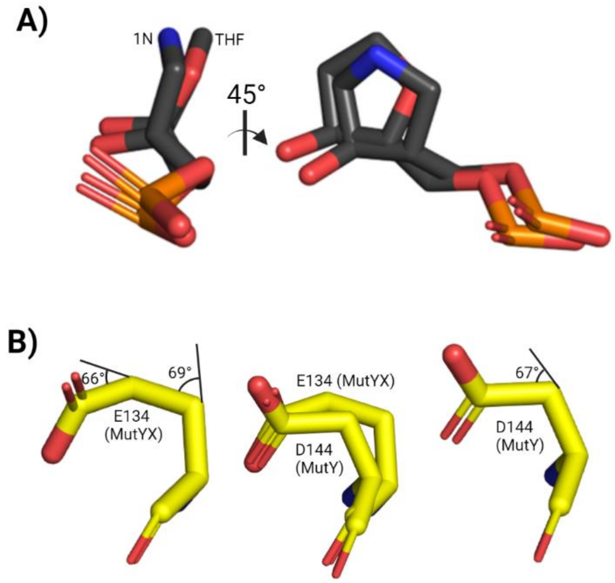

**Figure S5. A)** Structural alignment of the AP site analog (THF) from the MutYX structure and the transition state analog (1N) from GsMutY (PDB ID; 6UT7). **B)** structural comparison between the catalytic residues Glu134 and Asp144 from MutYX and GsMutY, respectively.

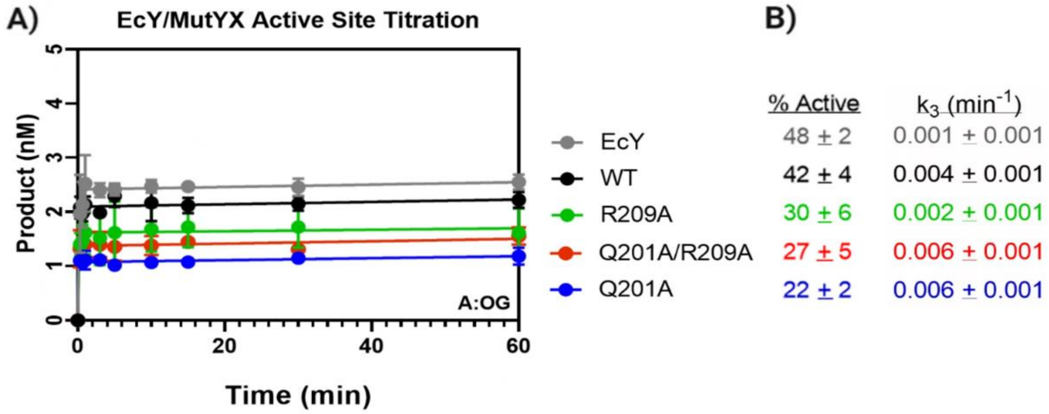

**Figure S6.** Multiple turnover (MTO) kinetics for determination of turnover rate, $k_3$ ($min^{-1}$) and active fraction (%) with OG:A substrate. **A)** Product production curves of MutYX or EcMutY enzymes under multiple turnover conditions, with 20 nM DNA substrate and ~5 nM enzyme concentration. **B)** Active fractions and rates of product release / turnover were obtained by assessing the amplitudes of MTO burst phase and slope of the linear phase, respectively.

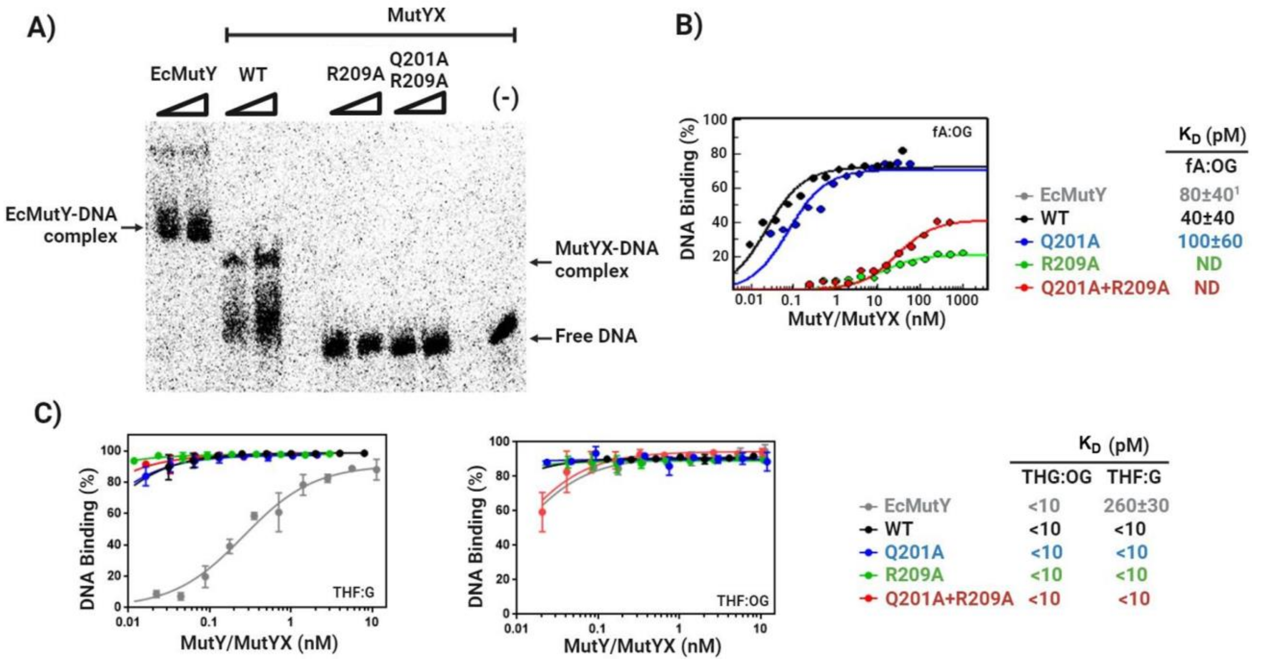

**Figure S7.** Electrophoretic mobility Shift Assays (EMSA) with WT and OG-recognition motif variants of MutYX. **A)** Qualitative EMSA using 0.5 µM and 1 µM enzyme showing that R209A and R209A + Q201A double mutant at the OG recognition sphere of MutYX do not form a shifted complex with the uncleavable substrate analog OG:fA DNA duplex. **B)** Plots for quantitation of percent bound as a function of [E] for MutY/MutYX with OG:fA. OG:fA-containing DNA (10 pM) was titrated with increasing concentration of enzyme (0.2 to 3 µM). The data was fit to a single-site binding isotherm to determine the dissociation constants ($K_D$) in the Table at the left. [1]$K_D$ value obtained from (3) **C)** Similar quantitative analysis for OG:THF and G:THF duplex, and $K_D$ determination.

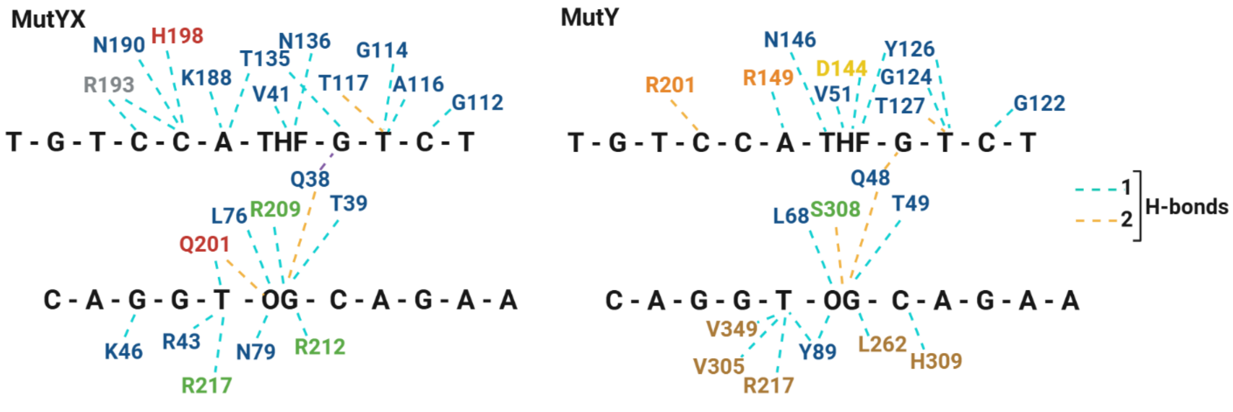

**Figure S8.** Scheme of DNA-protein contacts in GsMutY and MutYX structures. Residues from different domains or motifs are colored as follows: Catalytic domain, blue; X domain or FSH loop, green; IDC, red; Hw helix, gray; [4Fe-4S] cluster, orange; OG recognition domain, brown; and catalytic Asp, yellow.

| **Table S1**. Data Processing and Refinement Statistics for MutYX | |
|---|---|
| PDBID | 8UUC |
| Synchrotron (Beamline) | APS (24-ID-E) |
| Wavelength (Å) | 0.97918 |
| Space Group | $C222_1$ |
| Unit Cell Parameters | *a* = 38.83Å, *b* = 84.14Å, *c* = 225.74Å |
| Resolution Range (Å) | 40.0 – 1.55 (1.58 – 1.55) |
| No. observed reflections | 364,473 (18,204) |
| No. unique reflections | 53,001 (2,535) |
| Completeness | 97.6% (96.7%) |
| Multiplicity | 6.9 (7.2) |
| I/σ (I) | 12.8 (1.3) |
| $R_{merge}^{a}$ (%) | 6.3 (152.5) |
| $CC_{1/2}$ (%) | 99.9 (73.9) |
| **Refinement Statistics** | |
| $R_{factor}^{b}$ (%) | 17.6 |
| $R_{free}^{b}$ (%) | 19.8 |
| RMS bond length (Å) | 0.007 |
| RMS bond angle (°) | 1.034 |
| **Ramachandran Plot Statistics**[c] | |
| Favored (%) | 98.53 |
| Allowed (%) | 1.47 |
| Outliers (%) | 0.00 |
| **No. of atoms (Average B-factor)** | |
| Protein | 2,194 (33.5Å$^2$) |
| DNA | 439 (38.6Å$^2$) |
| $Na^+$ | 2 (40.1Å$^2$) |
| $Mg^{2+}$ | 2 (46.4Å$^2$) |
| $Cl^-$ | 1 (60.3Å$^2$) |
| Phosphate | 5 (1 molecule) (40.5Å$^2$) |
| Ethylene Glycol | 48 (12 molecules) (45.1Å$^2$) |
| Waters | 268 (41.7Å$^2$) |

[a] $R_{merge} = [\sum_h \sum_i |\mathbf{I}_h - \mathbf{I}_{hi}| / \sum_h \sum_i \mathbf{I}_{hi}]$ where $\mathbf{I}_h$ is the mean of $\mathbf{I}_{hi}$ observations of reflection h. Numbers in parenthesis represent highest resolution shell.

[b] R-Factor and [b] Rfree = $\sum||F_{obs}| - |F_{calc}|| / \sum|F_{obs}|$ x 100 for 95% of recorded data (R-Factor) or 5% data (Rfree).

[c] Ramachandran plot statistics from MolProbity (4).

**Table S2.** Results of the search of MMR components in *Eggerthella* genomes.

| Protein | Identified | Organism | Protein ID | GeneBank |
|---|---|---|---|---|
| MutS | No | | | |
| MutH | Possible | Eggerthella lenta | DUF559 domain-containing protein | WP_009305656.1 |
| | | Eggerthella lenta | YraN family protein | WP_015760617.1 |
| MutL | No | | | |
| SSB | Yes | Eggerthella sp. YY7918 | single-stranded DNA-binding protein | BAK43363.1 |
| RecJ | Yes | Eggerthella sp. YY7918 | single-stranded DNA-specific exonuclease | BAK45603.1 |
| ExoI | No | | | |
| UvrD | Yes | Eggerthella sp. YY7918 | helicase subunit of the DNA excision repair complex | BAK43813.1 |
| DNA Pol III | Yes | Eggerthella sp. YY7918 | DNA polymerase III | BAK44009.1 |
| DNA mismatch endonuclease Vsr | Yes | Eggerthella lenta DSM 2243 | DNA mismatch endonuclease Vsr | ACV56107.1 |
| G:T/U mismatch-specific DNA glycosylase | Yes | Eggerthella sp. YY7918 | DNA glycosylase | BAK44210.1 |

Note: MutS and MutL proteins were not identified in *Eggerthella* genomes. In DUF559 domain containing protein and YraN protein from *Eggerthela* were annotated as containing domains similar to MutH. However, based on amino acid sequence analysis it does not correspond to a MutH ortholog.
